# The nuclear transport receptor Impβ is a regulator of actin polymerization

**DOI:** 10.64898/2026.03.30.715331

**Authors:** Birthe Fahrenkrog, Hanmu Guo, Ahmed H. Mahmoud, Piotr Neumann, Philipp Armbruster, Chantal Rencurel, Richard W. Newton, Larisa E. Kapinos, Achim Dickmans, Roderick Y.H. Lim

## Abstract

Nuclear transport receptors are best known for mediating nucleocytoplasmic transport (NCT) through nuclear pore complexes. Here, we uncover an unexpected function of the primary import receptor importin-β (Impβ) as a direct regulator of the actin cytoskeleton. Impβ associates with stress fibers and the cell cortex and promotes actin polymerization through direct interactions. Disrupting Impβ-actin binding impairs stress fiber formation and suppresses cell migration well before NCT is affected. In 3D spheroids, perturbing Impβ further compromises tissue integrity, as reflected in changes to nuclear curvature and ellipticity. Together, these findings identify Impβ as a regulator of actin organization in both 2D and 3D contexts, revealing a direct link between the nuclear transport machinery and the cytoskeleton.

## Introduction

When cells encounter mechanical stresses, forces transmitted from focal adhesions via the cytoskeleton to the nuclear envelope can cause nuclear pore complexes (NPCs) to dilate or constrict (Lusk et al., 2025). These structural changes modulate nucleocytoplasmic transport (NCT) of mechanosensitive transcription factors such as Yes-associated protein (YAP) (Elosegui-Artola et al., 2017). In this way, mechanical cues are converted into biochemical signals and subsequently gene expression programs (Uhler and Shivashankar, 2017), which is central to understanding how cells regulate migration, morphogenesis and tissue organization (Iskratsch et al., 2014; Kalukula et al., 2022; Wang et al., 2009b; Wozniak and Chen, 2009). However, this form of “outside-in” mechanotransduction implicitly treats NCT as a passive responder to force and whether the NCT machinery actively participates in mechanotransduction to establish homeostatic feedback from the “inside-out” remains unknown. This issue is of fundamental interest as mechanical abnormalities in cells are associated with diseases such as cancer (Ingber, 2003; Isermann and Lammerding, 2013; Plodinec et al., 2012).

Nuclear transport receptors of the karyopherin-β family mediate NCT by selectively shuttling signal-bearing cargoes through NPCs (Wing et al., 2022). These receptors include importins and exportins that partition asymmetrically in the cytoplasm and nucleus, respectively (Kalita et al., 2021; Wuhr et al., 2015). Although its origin remains unclear, this asymmetry likely supports the directional transport of cargo, with importins carrying nuclear localization signal (NLS)-cargoes into the nucleus and exportins ushering nuclear export signals (NES)-cargoes out of it (Fried and Kutay, 2003; Kehlenbach et al., 2023). Inside the nucleus, the guanosine triphosphate-bound form of GTPase Ran (RanGTP) releases NLS-cargo from importins and promotes assembly of NES-cargo-exportin-RanGTP complexes for export (Fornerod et al., 1997). RanGTP is then hydrolyzed to RanGDP in the cytoplasm, which frees importins and exportins for subsequent rounds of NCT.

Beyond nucleocytoplasmic transport, the primary import receptor Impβ (Impβ or Kapβ1; (Gorlich et al., 1995a; Gorlich et al., 1995b; Imamoto et al., 1995a; Imamoto et al., 1995b; Moroianu et al., 1995)) is recognized for its functional versatility (Harel and Forbes, 2004). As a cytoplasmic chaperone, Impβ protects basic proteins against aggregation as implicated in neurodegenerative disorders (Doll and Cingolani, 2022; Guo et al., 2018; Hayes et al., 2020; Jakel et al., 2002; McGoldrick et al., 2023). Impβ controls spindle formation and positioning during mitosis by binding spindle assembly factors and preventing their interaction with microtubules in a Ran-dependent manner (Ciciarello et al., 2004; Ems-McClung et al., 2004; Gruss et al., 2001; Schatz et al., 2003; Wiese et al., 2001). Impβ-Ran signaling further contributes to microtubule-based transport in neurons (Alber et al., 2023; Baameur et al., 2016) and coordinates NE and NPC reassembly after mitosis (Askjaer et al., 2002; Bamba et al., 2002; Hachet et al., 2004; Harel et al., 2003; Hetzer et al., 2000; Hetzer et al., 2001; Ryan et al., 2003; Timinszky et al., 2002; Walther et al., 2003; Zhang and Clarke, 2000; Zhang and Clarke, 2001). Interestingly, Impβ also interfaces with actin-based systems. F-actin polymerization and cytoskeletal tension control the nuclear import of Myocardin-related transcription factors (MRTFs), linking Impβ and actin dynamics (Mouilleron et al., 2011). Likewise, actin-based scaffolds at the cell periphery regulate the accessibility of signaling transcription factors such as YAP, NF-κB, and MAP kinases to importins (Aplin and Juliano, 2001; Campbell and Hope, 2003; Cyert, 2001). Impβ signaling is also implicated in actin-dependent processes such as cortical actomyosin dynamics during cytokinesis and actin-remodeling in endothelial inflammation (Baameur et al., 2016; Leonard et al., 2018; Ozugergin and Piekny, 2021; Samwer et al., 2013).

These observations suggest that Impβ interfaces with actin networks, yet whether Impβ directly regulates actin dynamics remains unexplored. Here, we show that Impβ interacts physically and functionally with the actin cytoskeleton, separate from its role in NCT. Impβ binds both monomeric and filamentous actin and enhances actin polymerization. Interfering with Impβ function/activity disrupts F-actin organization, leading to altered cell and nuclear morphology, impaired cell migration, and compromised integrity of 3D cell culture spheroids. Furthermore, importazole (IPZ) (Soderholm et al., 2011), an inhibitor of the Impβ import pathway, also hinders Impβ-actin binding and rapidly collapses the actin cytoskeleton within minutes, whereas disruption of NCT takes hours. These findings identify Impβ as a direct regulator of actin dynamics, thereby linking NCT to the mechanotransduction apparatus.

## Results

### Impβ associates with the cell cortex and actin stress fibers

Ori 3.1 (also referred to as Nthy-ori 3-1 cells) is a normal thyroid follicular epithelial cell line immortalized by SV-40 large T antigen transfection (Landa and Robledo, 2011; Zhao et al., 2011). Unlike conventional nuclear transport assays that image the nuclear mid-plane, all images in this study were acquired along the basal plane (unless stated otherwise) where the cytoskeleton is most prominent (Fig. 1A). Immunofluorescence combined with confocal imaging revealed Impβ enrichment at the cell cortex and leading edge (Fig. 1B, panel 1), in addition to its known localization in the nucleus, cytoplasm, and at NPCs. Consistent with basal plane imaging, NPCs appear as evenly distributed puncta across the nuclear surface (Dultz and Ellenberg, 2010), rather than the typical punctate rim-like staining observed near the mid-plane (Sakuma et al., 2025). Nano-secondary antibodies or fluorophore-conjugated primary antibodies revealed a more highly resolved association of Impβ with filamentous cytoplasmic structures resembling actin stress fibers (Fig. 1B, panel 2-4). Likewise, Ori 3.1 cells expressing endogenous mClover-tagged Impβ (Impβ-mClover) or exogenous enhanced green-fluorescence protein (EGFP) tagged Impβ (Impβ-EGFP) localized to the cell cortex and to stress fiber-like structures (Fig. 1C; Supplemental Movie 1). Phalloidin staining (Estes et al., 1981; Pospich et al., 2020; Wieland and Govindan, 1974) confirmed the association of Impβ with actin in Ori 3.1 cells (Fig. 1D, upper row) and in Ori 3.1 Impβ-mClover cells (Fig. 1D, lower row). Digitonin permeabilization markedly improved visualization of Impβ along actin stress fibers (Fig. 1E) by removing cytoplasmic proteins and reducing background fluorescence (Adam et al., 1990).

**Figure 1:**
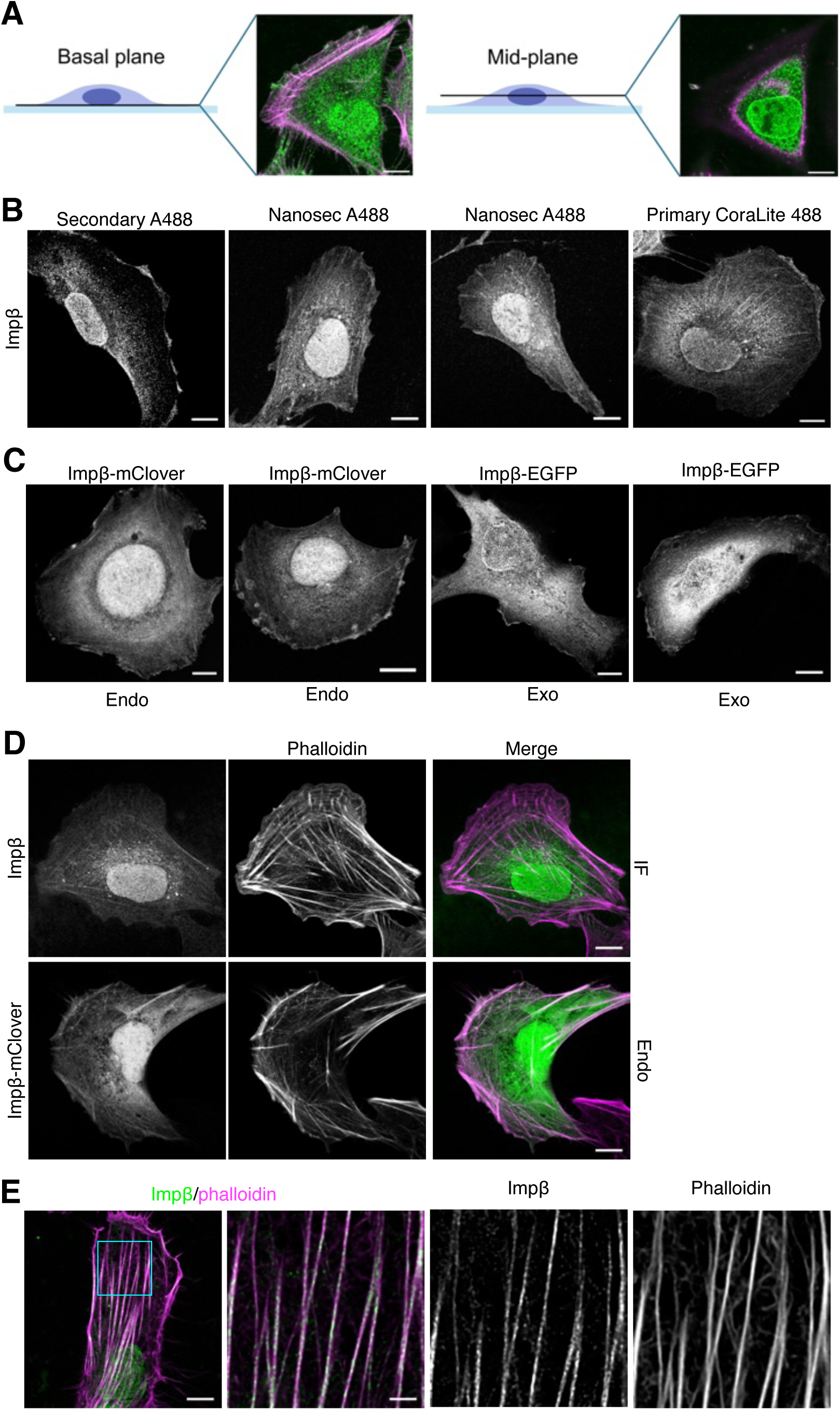
Impβ localizes to the cell cortex and actin stress fibers. (**A**) Schematic summary depicting imaging positions in cells cultured on glass. (**B**) Ori 3.1 thyroid epithelial cells were immunostained for Impβ (Impβ) using secondary Alexa 488 conjugated antibodies (secondary A488; left), Nano-secondary Alexa 488 conjugated secondary antibodies (Nanosec A488; middle 2), or CoraLite488 conjugated primary antibody (Primary CoraLite 488; right). Impβ localizes to the nucleus, the cytoplasm, and nuclear envelope (NE). At higher resolution using Nano-secondary antibodies or directly conjugated primary antibodies an association with actin stress-fibers becomes evident. (**C**) Endogenous importin-β-mClover (Impβ-mClover) and exogenous importin-β-EGFP (Impβ-EGFP) similarly associate with actin stress fibers, besides localizing to the nucleus, the cytoplasm, and NE. (**D**) To confirm the association of Impβ with actin, Ori 3.1 cells as well as Impβ-mClover expressing cells were co-stained for Impβ and phalloidin to visualize actin stress fibers. Shown are representative confocal images and (**E**) deconvoluted confocal images, of either Triton-X100 (upper row) or digitonin (lower row) permeabilized Ori 3.1 cells. Scale bars, (**A**-**D**, 10 µm; **D**, zoom in 2.5 µm).

To ascertain the generality of such behavior, we found that Impβ also associated with the cell cortex and actin stress fibers in MRC5 fibroblasts (Fig. S1A), as well as cell lines originating from different tissues and (patho)physiological states. These included the fibroblast AG08468 cell line, ARPE-19 cells from spontaneously immortalized retinal pigment epithelia, as well as MDA-MB231 triple negative breast cancer cells, and cervical cancer HeLa cells (Fig. S1B). Furthermore, the association of Impβ with the cell cortex appears independent of whether the cells were grown on glass or on collagen IV-coated glass coverslips (Fig. S1C). Indeed, their association persisted even after filamentous actin was disrupted using RhoA/ROCK kinase inhibitor Y27632 (Ishizaki et al., 2000), cytochalasin B (Shoji et al., 2012) and latrunculin A (Ayscough et al., 1997; Coue et al., 1987; Fujiwara et al., 2018) (Fig. S1D), indicating that the association of Impβ with actin is not limited to stress fibers.

### Impβ interacts with actin

Actin has been identified as a potential Impβ interactor in two proteomic studies (Di Francesco et al., 2018; Song et al., 2022). To establish their interaction in cells, we used bimolecular fluorescence complementation (BiFC), which reports protein-protein interactions through fluorescence reconstitution of split fluorescent protein fragments (Kodama and Hu, 2010). Human Impβ comprises 19 HEAT (**H**untingtin, **E**longation factor 3, protein phosphatase 2**A**, **T**arget of rapamycin (TOR)) repeats (Andrade et al., 2001), structurally arranged in two nearly anti-parallel α-helices, termed A- and B-helices, connected by short linkers (Cook et al., 2007). Moreover, a splice variant lacking the first 145 residues (isoform 2) might exist in cells (Fig. 2A; www.ncbi.nlm.nih.gov/gene/3837) (Ota et al., 2004). Split-Venus protein constructs, comprising N-terminal (V1-(or VN-)) fragments were used to generate V1 fusion proteins with full-length Impβ (residues 1-876), isoform 2 (Iso2, residue 146-876), the N-terminal region (N145, residues 1-145), and HEAT repeat 1 (N31, residues 1-31) (Fig. 2A). Meanwhile, actin was fused to a C-terminal (V2- (or VC-)) fragment.

**Figure 2:**
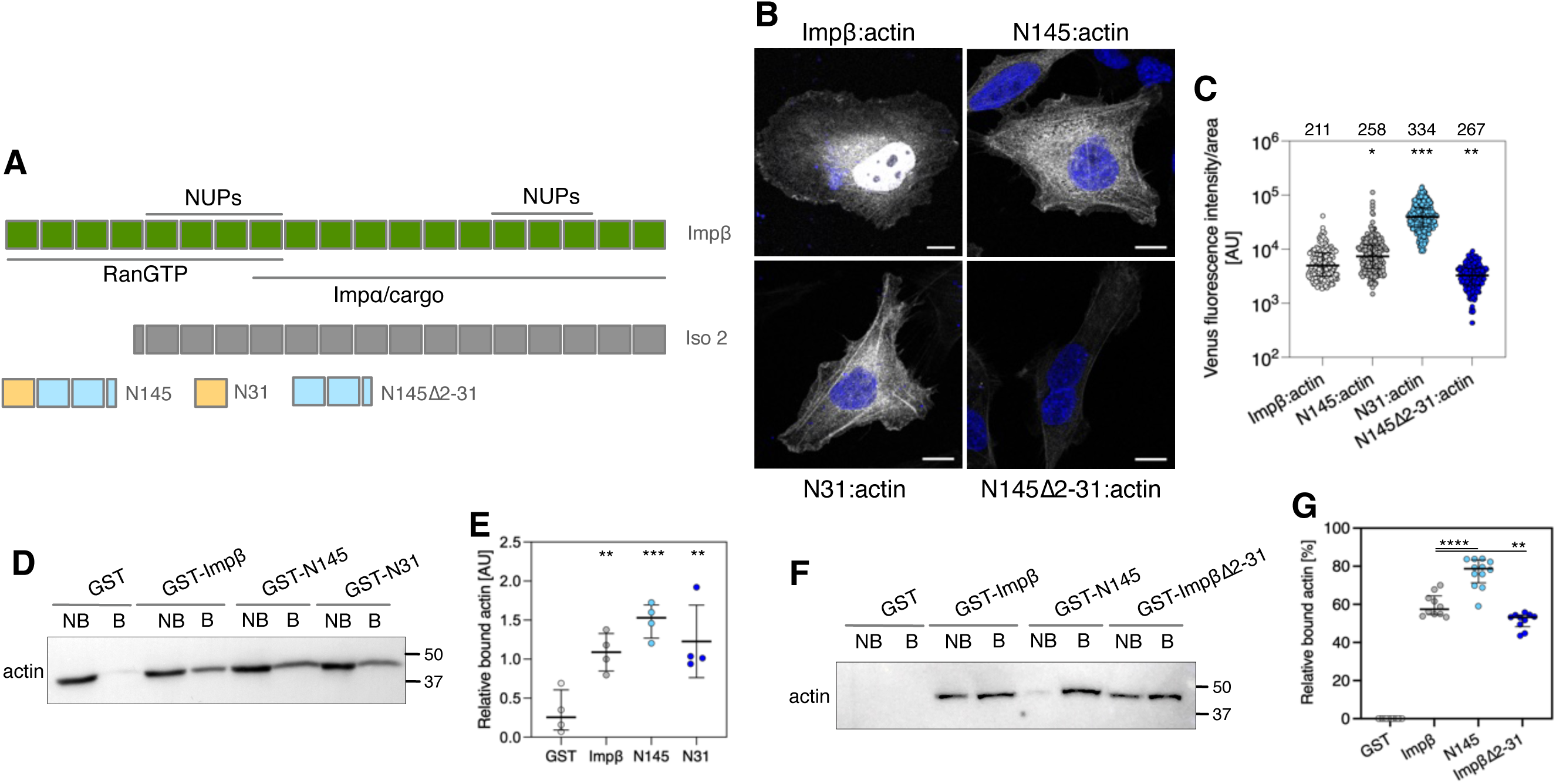
Identification of actin as an Impβ binding partner. (**A**) Impβ (Impβ) fragments utilized in this study with binding regions to different molecular partners indicated. Regions to different binding partners are indicated. (**B**) Bimolecular fluorescence complementation (BiFC) signal produced by interaction between Impβ and actin. Actin interacts with full-length Impβ (Impβ:actin), the N terminal 145 residues (N145:actin), and the N terminal 31 residues (N31:actin). Deletion of residues 2-31 (i.e., HEAT repeat 1; N145Δ2-31:actin) largely abolished the interaction of Impβ with actin. DNA was stained with DAPI (blue). Shown are representative images from at least three independent experiments. (**C**) Quantitative analysis of the BiFC signals for each condition. Black horizontal line represents the median with interquartile range. The number of analyzed cells is indicated at the top of each column. Two-Way Anova test was used to calculate statistics: ***p <0.001, **p <0.01, *p <0.05. (**D**) GST pull down-assays were performed using recombinant Impβ fragments fused to GST (GST-Impβ, GST-N145, and GST-N31) and GST alone. Fragments were immobilized on magnetic glutathione-agarose beads and incubated with Ori 3.1 cell lysates. Bound and non-bound fractions were analyzed by immunoblotting using anti-actin antibodies and (**E**) quantified. (**F**) *In vitro* binding assays were run using the indicated recombinant GST- Impβ fragments immobilized on magnetic glutathione-agarose beads in combination with recombinant human β-actin. Bound and non-bound fractions were analyzed by immunoblotting using anti-actin antibodies and (**G**) quantified. Band intensities of the bound fractions were normalized to the respective non-bound fraction. Values are the median with interquartile range. Two-way Anova test, ****p <0.0001 ***p <0.001, **p <0.01. N=4 (independent pull down as well as *in vitro* binding assays).

Expressing Impβ-V1, N145-V1, and N31-V1 together with actin-V2 in Ori 3.1 cells resulted in strong fluorescence in the cytoplasm, the cell cortex and along actin stress fibers (Fig. 2B). Impβ:actin complexes also produced nuclear fluorescence. Since BiFC complexes are irreversible (Kodama and Hu, 2010), Impβ may retain partial import activity in the BiFC complex with actin. In contrast, the N145:actin and the N31:actin complexes lack import competence (Fig. 2A) and are therefore predominantly cytoplasmic, with only a minor fraction entering the nucleus by passive diffusion (Fig. S2A). However, deleting HEAT repeat 1 (N145Δ2-31) dramatically decreased the Venus fluorescence signal, suggesting that HEAT repeat 1 is essential for Impβ-actin binding. To facilitate quantification, experiments were repeated in HeLa cells (Fig. S2A), which have a higher transfection efficiency than Ori 3.1 cells. HeLa cells were imaged with constant confocal laser power to enable quantitative comparison of fluorescence signals across cells (Fig. 2C). This analysis revealed that N31:actin and N145:actin produced significantly stronger signals than Impβ:actin, whereas the N145Δ2-31:actin signal was significantly lower. In contrast, no fluorescence signals were detected for isoform 2:actin, Impβ:Arp2 or Impβ:β-tubulin thereby confirming the specificity of the Impβ:actin interaction (Fig. S2B). Likewise, Impα:actin, Impβ2:actin, XPO1:actin, and Ran:actin pairs did not generate a fluorescence signal (Fig. S2C), whereas known interacting pairs of Impβ:Impα, actin:vinculin, and actin:myosin confirmed the validity of the method (Fig. S2D). Together these data indicate that Impβ specifically binds actin and that HEAT repeat 1 (residues 1-31) is necessary and sufficient for this interaction in cells. In line with this, Impβ-EGFP, N145-EGFP, and N31-EGFP also associated with the cell cortex and actin filaments in Ori 3.1 and HeLa cells, whereas Impβ-Δ2-31-EGFP and Iso2-EGFP did not (Fig. S2E).

To confirm the role of HEAT repeat 1 in mediating the Impβ-actin interaction, recombinantly expressed GST-fusion proteins of full-length and truncated Impβ were immobilized on magnetic glutathione-beads and incubated with Ori 3.1 cell lysates or purified recombinant monomeric (G-)human β-actin. Western blot analysis of bound and non-bound fractions revealed that GST-Impβ, GST-N145, and GST-N31 pulled-down significantly more actin from cell lysates than GST alone (Fig. 2D and E). Moreover, human β-actin bound GST-Impβ and GST-N145, whereas the amount bound to GST-ImpβΔ2-31 (deletion of HEAT repeat 1) was significantly diminished (Fig. 2F and G). These assays confirm that Impβ binds actin with HEAT repeat 1 making a substantial contribution to this interaction.

We then used Alphafold2 (https://alphafold.ebi.ac.uk; (Jumper et al., 2021)) to predict residues in Impβ HEAT repeat 1 and actin that mediate their interaction. Although the aligned error (PAE) scores were relatively low, the PDB interface analysis (PISA) identified close contacts between Impβ E8 and actin S350 (3.09 Å) and between Impβ Q22 and actin E167 (3.36 Å) (Fig. 3A). Targeted mutations of these residues in either N31-V1 (N31-E8AQ22A) or actin-V2 (actin-E167AS350A) significantly reduced BiFC fluorescence. Co-expression of both mutant fragments diminished the signal even further (Fig. 3B and C). Pairwise mutations to N31 and actin (E8A and S350A; Q22A and E167A) likewise compromised the Venus signal (Fig. S3B). These effects were also reproduced in HeLa cells (Fig. S3C) as well as in Ori 3.1 cells expressing full-length Impβ (Fig. S3D). Collectively, these results indicate that HEAT repeat 1 plays a key role in mediating Impβ-actin interactions.

**Figure 3:**
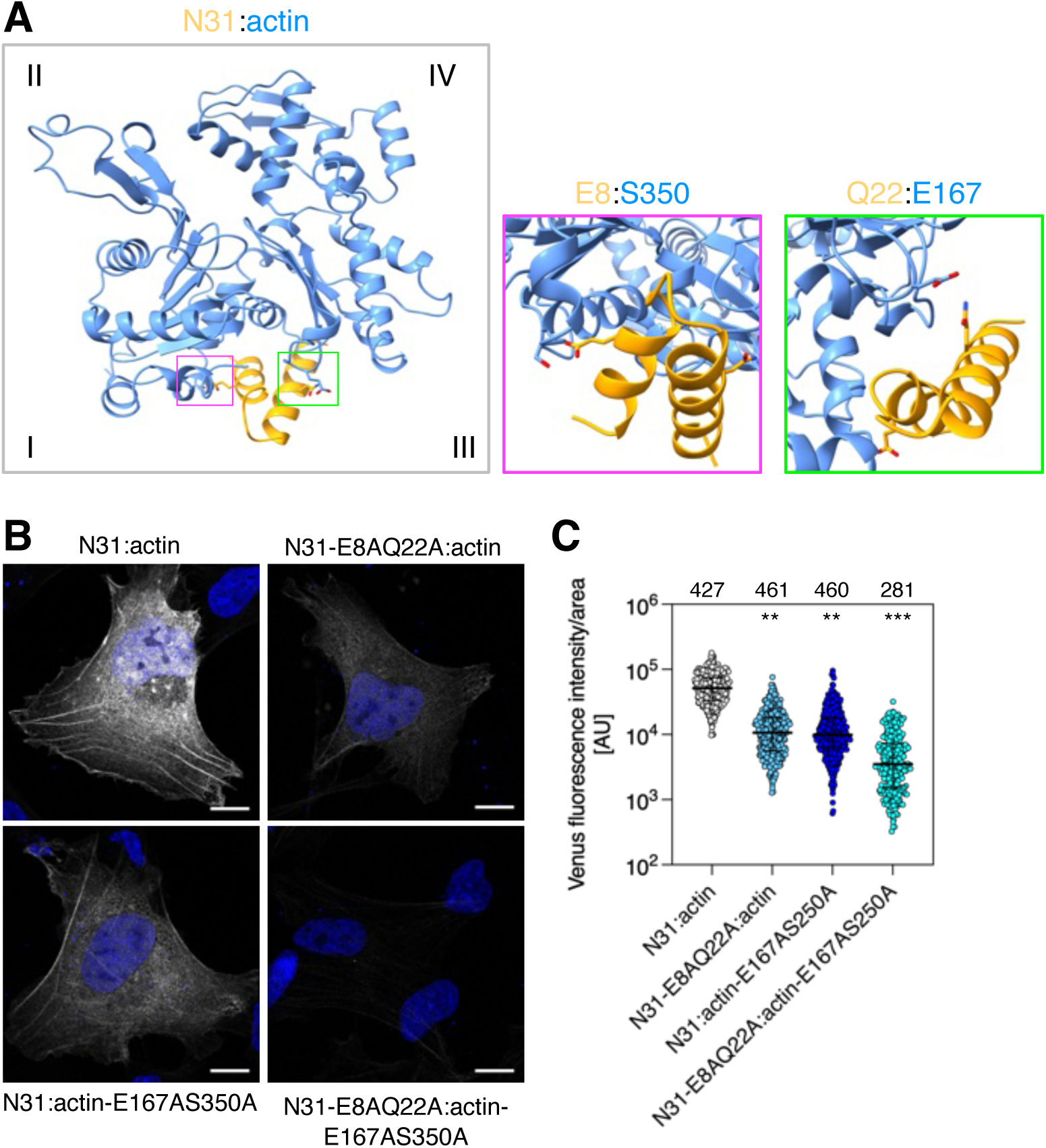
Residues in HEAT repeat 1 of Impβ are implicated in the interaction with actin. (**A**) AlphaFold2 structural prediction of HEAT repeat 1 binding to actin. Residues E8 and Q22 of Impβ likely bind to S350 and E167, respectively, of actin. The actin subdomains are indicated by roman numerals. (**B**) Interaction of N31:actin shown by BiFC. Mutating E8 and Q22 of Impβ to alanine (N31-E8AQ22A) and S350 and E167 of actin (actin-E167AS250A) largely reduced the interaction of Impβ with actin, while mutations in all four residues abolished the interaction. DNA was stained with DAPI (blue). Shown are representative images from at least three independent experiments. Scale bars, 10 µm. (**C**) Quantitative analysis of the BiFC signals for each condition. Black horizontal line represents the median with interquartile range. The number of analyzed cells is indicated at the top of each column. Two-Way Anova test was used to calculate statistics: **, p < 0.01; ***, p <0.001.

### Importazole inhibits the Impβ-actin interaction

Importazole (IPZ) inhibits the RanGTP–Impβ interaction but its mechanism remains unclear(Jans et al., 2019; Soderholm et al., 2011; Vercruysse et al., 2024). However, because Ran binds in the N-terminal region of Impβ (PMID: 10367892, PMID: 15864302), we hypothesized that IPZ may also perturb the Impβ-actin binding interface. Subsequent AlphaFold3 structural modelling of the Impβ-IPZ interface supported this notion and indicated that the binding site of IPZ on Impβ is in close proximity to the potential actin binding site, namely to Phe24 (Fig. S4A). Using surface plasmon resonance (SPR), we found that IPZ binds both full-length Impβ and its N145 fragment with KD values of about 37 µM for Impβ and 41 µM, respectively (Fig. S4B), which is consistent with the concentration required to inhibit nuclear import (Soderholm et al., 2011). Importantly, this indicates that the N-terminal region of Impβ is the primary binding site for IPZ.

### Importazole binding to Impβ perturbs the interaction with actin

We then examined Impβ binding to G- and F-actin *in vitro* using high speed (150.000 x g) co-sedimentation assays. Adding recombinant Impβ to G-actin enhanced actin polymerization and kinetics in a concentration-dependent manner, comparable to the actin nucleator mDia2 (Fig. 4A-F). While ∼4 % of Impβ sedimented in the absence of actin (control), ∼25% co-sedimented with F-actin, indicating a direct interaction that was further supported by negative-staining electron microscopy and dual-color stimulated emission depletion (STED) imaging (Fig. 4G-K). Pre-incubating Impβ with IPZ reduced F-actin binding (Fig. 5A), lowering Impβ co-sedimentation to ∼10% and reduced F-actin density (Fig. 5B-D). IPZ alone had no effect on filament density, integrity, or actin sedimentation (Fig. 5A-D; Fig. S4C and D). Together, these assays confirmed that Impβ inhibition by IPZ negatively affects actin filament organization.

**Figure 4:**
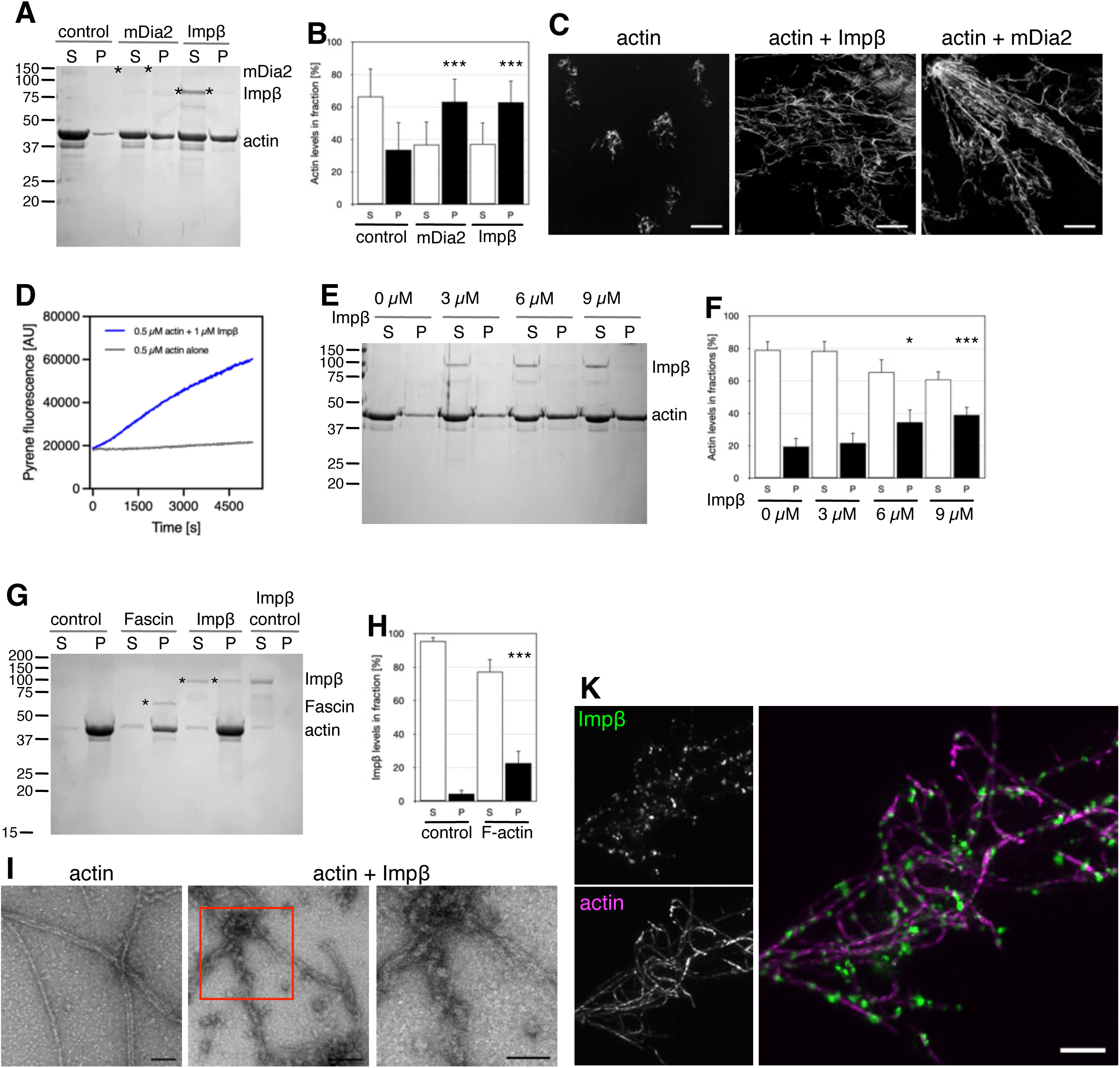
Impβ enhances actin polymerization and binds F-actin. (**A**) SDS-PAGE analysis (12% acrylamide) of the effect Impβ on actin polymerization. G-actin (rabbit, skeletal muscle) was co-incubated with 6 µM Impβ for 30 minutes prior polymerization and high-speed centrifugation. The supernatant (S) and pelleted (P) fraction were analyzed by SDS-PAGE. mDia2 served as control for a protein enhancing actin polymerization. (**B**) Quantification of actin levels in the supernatant and pelleted fractions. Number of assays: N=6-7. Values are means ± s.d. *, p < 0.05, ***, p < 0.001, T-test, two-tailed. (**C**) Confocal imaging of the polymerization products outlined in (**A**). Actin filaments were stained with phalloidin-iFluor594 and mounted on glass using Mowiol. Scale bars, 10 µm. (**D**) Polymerization kinetics of 0.5 µM monomeric actin (5% pyrene-labeled) in the presence of 1 µM Impβ. (**E**) G-actin was co-incubated with the indicated increasing amounts of Impβ for 30 minutes before polymerization and high-speed centrifugation. The supernatant (S) and pelleted (P) fractions were analyzed by 12% SDS-PAGE. (**F**) Quantification of actin levels in the supernatant and pelleted fraction. Number of assays: N=5. Values are means ±s.d. *, p < 0.05, ***, p < 0.001; T-test, two-tailed. (**G**) F-actin was co-incubated with 6 µM Impβ for 30 minutes and sedimented by high-speed centrifugation. The supernatant (S) and pelleted (P) fractions were analyzed by 12% SDS-PAGE. Fascin served as control for a F-actin-binding protein. (**H**) Quantification of Impβ and actin levels in the supernatant and pelleted fractions. Number of assays: N=3-5. Values are means ±s.d. ***, p < 0.001; T-test, two-tailed. (**I**) EM micrographs of the reaction products as outlined in (**F**). Actin filaments were stabilized with 6 µM phalloidin and subjected to negative staining EM. Scale bars, 100 nm (left and middle), 50 nm (right). (**K**) Stimulated emission depletion (STED) imaging of phalloidin-iFluor594 labelled F-actin (magenta) and Impβ stained with Abberior Star635P. Scale bar, 2 µm.

**Figure 5:**
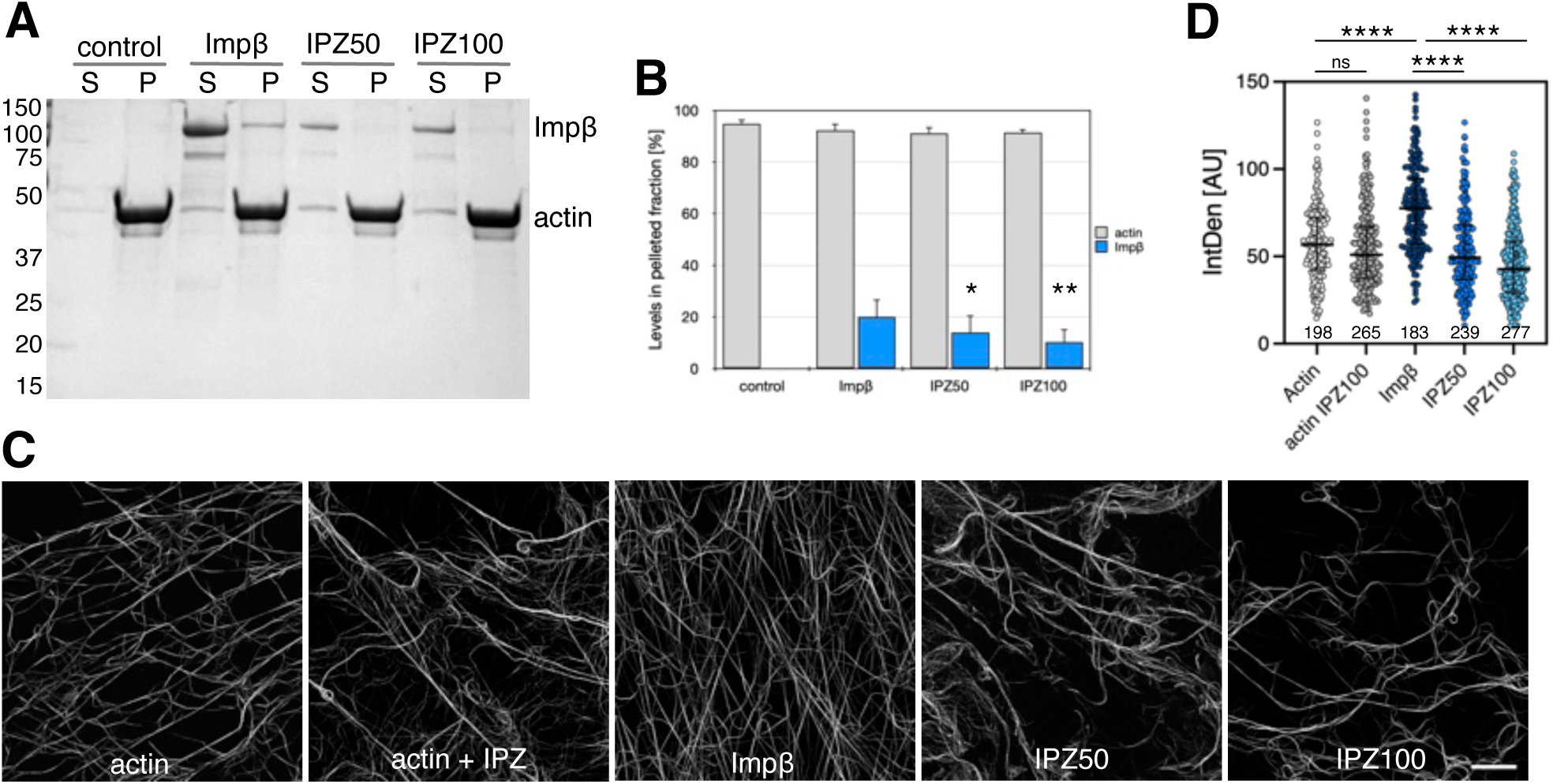
Importazole compromises Impβ binding to actin. (**A**) SDS-PAGE analysis (12% acrylamide) on the effect of pre-incubation of Impβ with importazole (IPZ). Impβ was incubated for 30 minutes with 50 µM IPZ (IPZ50) and 100 µM IPZ (IPZ100) prior to incubation with F-actin. The supernatant (S) and pelleted (P) fraction after high-speed centrifugation were analyzed by SDS-PAGE and (**B**) the respective levels of Impβ and actin in pelleted fractions were quantified. Number of assays: N=4. Values are means ± s.d., *p<0.05, **, p < 0.01, T-test, two-tailed. (**C**) Confocal imaging of the polymerization products outlined in (**A**). Actin filaments were stained with phalloidin-iFluor594 and mounted on glass using Mowiol. Scale bar, 10 µm. (**D**) Quantification of the F-actin density (IntDen) of the polymerization products. Welch’s t-test (unpaired) was used to calculate statistics: **, p < 0.01, ****, p < 0.0001.

### Impβ inhibition disrupts actin organization and alters cell morphology

With this evidence in hand, we treated Ori 3.1 cells with IPZ to determine whether Impβ-actin binding could be inhibited in cells. Indeed, IPZ treatment markedly reduced the number of stress fibers (Fig. 6A) compared with control cells or cells treated with ivermectin (IVM), which inhibits importin-α (Impα; (Jans et al., 2019; Martin and Jans, 2021)). Quantitative analysis using FilamentSensor 2.0 (Fig. S5A) (Hauke et al., 2023) confirmed that IPZ significantly decreased the number of actin filaments, which were also shorter and thicker (Fig. 6B-D). Similarly, depleting Impβ by siRNA affected length and width of actin filaments, whereas the overall number was comparable to cells treated with non-targeting siRNAs (Fig. 6E-G). On the contrary, depleting Impα (siKPNA2) or the nuclear export receptor CRM1/XPO1 (siXPO1; Fig. 6E-G) did not affect actin filaments. Cytoskeleton integrity is required for the mechanical regulation of YAP import into the nucleus (Das et al., 2016; Elosegui-Artola et al., 2017; Piccolo et al., 2014). Accordingly, IPZ treatment markedly reduced nuclear YAP compared with control or IVM treated cells, consistent with disruption of the actin cytoskeleton (Fig. 6H and I).

**Figure 6:**
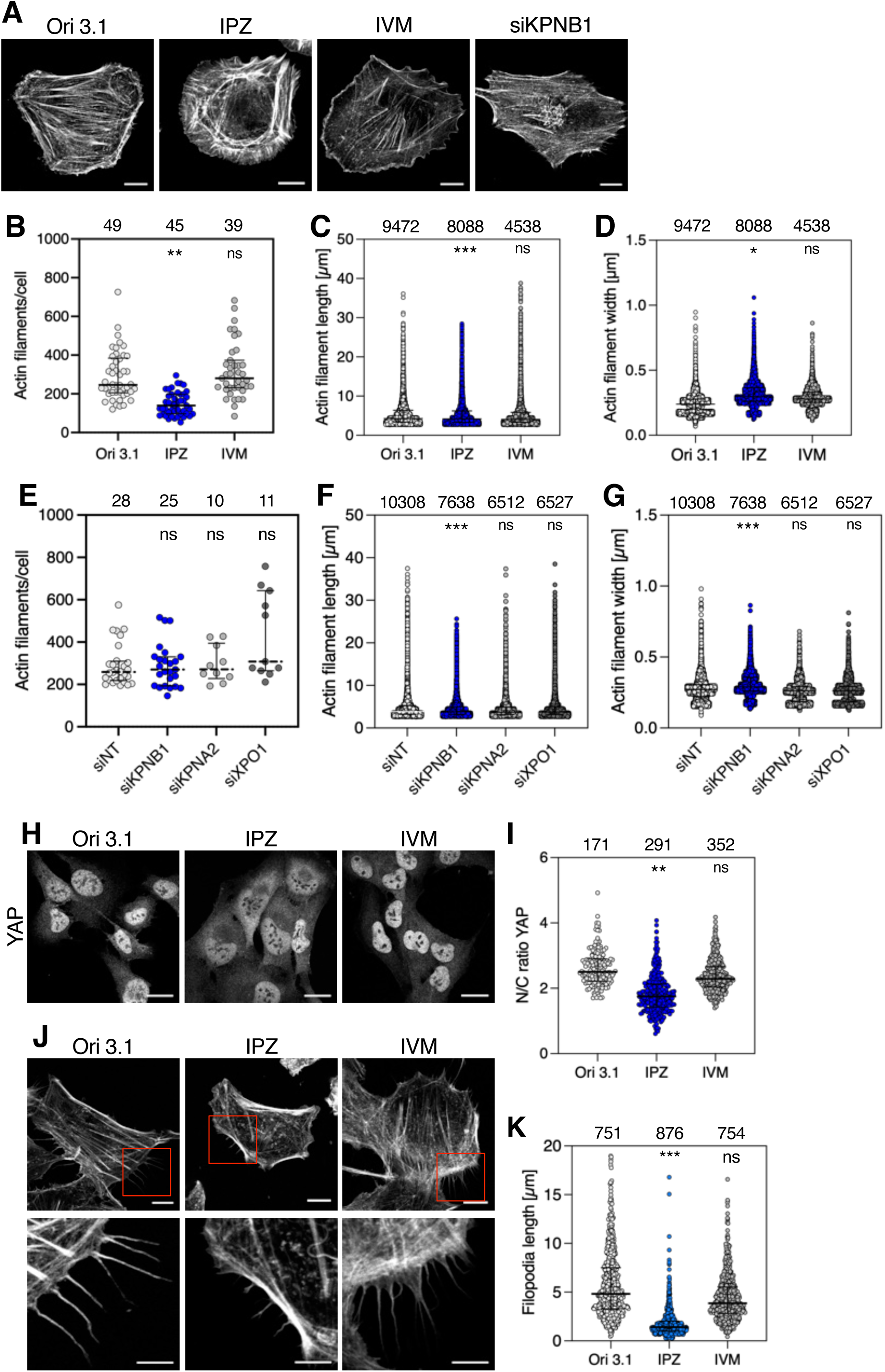
Inhibition/depletion of *Impβ* affects actin stress fiber and filopodia formation. Ori 3.1 cells were grown on glass cover-slips and treated with 50 µM importazole (IPZ) or 2.5 µM ivermectin (IVM) for 1 h or siRNAs against Impβ (siKPNB1) for 48 h. Cells were stained with phalloidin to visualize F-actin. (**A**) Confocal images were recorded (upper row) and actin stress fibers were revealed (Fig. S5) using the open-source JRE analysis tool FilamentSensor II. Quantitative analysis of actin stress fibers summarizing (**B, E**) the number of actin filaments per cell, as well as (**C, F**) the length, and (**D, G**) the width of actin filaments. Black horizontal line represents the median with interquartile range. The number of analyzed cells and filaments, respectively, is indicated at the top of each column. Two-Way Anova test was used to calculate statistics: *, p < 0.05, **, p < 0.01, ***, p < 0.001; ns, non-significant. (**H**) IPZ, but not IVM treatment of Ori 3.1 cells perturbs YAP1 localization in the nucleus. Shown are confocal images of YAP1-stained cells. The YAP nuclear-cytoplasmic ratio (N/C ratio) was measured using the Fiji/ImageJ plug-in “Nuclei_cytoplasm_measuring” and quantified (**I**). The number of analyzed cells and filaments, respectively, is indicated at the top of each column. Black horizontal line represents the median with interquartile range. Two-Way Anova test was used to calculate statistics: **, p < 0.01; ns, non-significant. (**J**) IPZ, but not IVM treatment of Ori 3.1 cells affects filopodia formation. Shown are confocal images of phalloidin-stained cells. Filopodia length was measured using Fiji/ImageJ and quantified (**K**).

Similarly, treating Ori 3.1 Impβ-mClover cells with IPZ reduced both the number and length of actin filaments (Fig. S5B-E), as well as the filamentous “imprints” formed by Impβ-mClover (Fig. S5B, F-H). Notably, these effects were detectable within just 5 minutes of IPZ treatment (Fig. S5B-H). At this early time point, however, neither nuclear import of 53BP1 (Fig. S6A and C), an Impα/Impβ-dependent cargo (Matsuura, 2019; Moudry et al., 2012; von Morgen et al., 2018), nor the intracellular distribution of Impβ itself was affected (Fig. S6B and D). This is consistent with reports showing that nuclear import defects become detectable only after 6 hours of IPZ treatment (Ge et al., 2019; Kublun et al., 2014). These observations highlight the rapid mechanosensitivity of actin assembly and disassembly compared with the slower, diffusion-driven kinetics of NCT (Wang et al., 2009b). IPZ treatment also produced significantly shorter filopodia than in control and IVM-treated cells (Fig. 6J and K).

Thereafter, we cultured Ori 3.1 cells as spheroids to study Impβ inhibition in a 3D model system, which provides a closer approximation of *in vivo* conditions. Imaging of the spheroid mid-plane showed that IPZ treatment disrupted the “tensional skin”, a thin, high-tensioned outer layer of cells that counterbalances internal compressive stresses (Kosheleva et al., 2023; Lee et al., 2019; Riccobelli, 2025), leading to reduced spheroid compactness (Fig. 7A-C). This coincided with reduced nuclear ellipticity in cells at the spheroid periphery (Fig. 7C and D), whereas nuclei near the spheroid center were unaffected (Fig. 7A and E). In contrast, inhibition of Impα by IVM had no detectable effect on spheroid integrity or nuclear ellipticity (Fig. 7C and D). Together, these findings show that Impβ inhibition rapidly disrupts actin organization and cellular mechanics, consistent with a direct, NCT-independent role in regulating actin dynamics.

**Figure 7:**
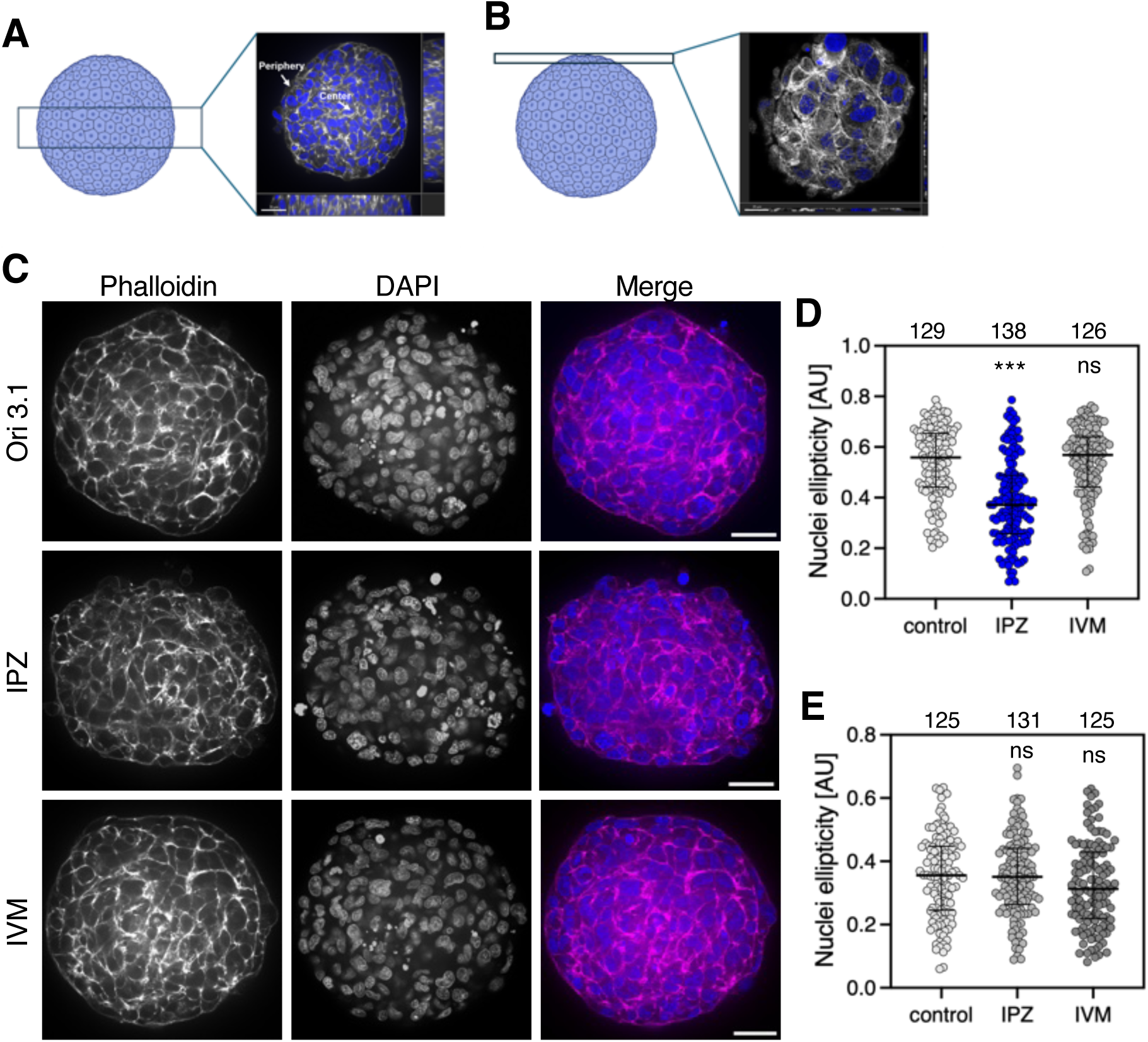
Inhibition of Impβ perturbs Ori 3.1 spheroids integrity. **(A, B)** Schematic summary depicting imaging positions in spheroids. (**C**) Ori 3.1 cells were grown on ultralow binding plates for 3 days and treated with 75 µM IPZ or 2.5 µM IVM for 3 hours prior to staining with phalloidin to visualize F-actin (magenta). DNA was stained with DAPI (blue). Shown are representative spinning disc confocal images of peripheral cells of spheroids. Scale bars, 40 µm. (**D**) The ellipticity of nuclei in the peripheral and (**E**) central cells of spheroids were quantified using Fiji/ImageJ (see also Fig. S7A). A total number of 20 spheroids from 4 independent experiments were analyzed. The number of analyzed cells is indicated at the top of the graph. Dunn’s multiple comparison/Kruskal-Walllis test was used to calculate statistics: ***, p < 0.001; ns, non-significant. (**F**) Quantitative analysis of actin filament (Fig. S7B) density, (**G**) length, and (**H**) width using the open-source JRE analysis tool FilamentSensor II. Black horizontal line represents the median with interquartile range. The number of respective analyzed cells and filaments are indicated at the top of each column. Two-way Anova test was used to calculate statistics: ****, p < 0.0001, ns, non-significant.

### Inhibition of Impβ affects cell migration

Finally, we used wound healing assays to assess whether inhibiting Impβ function affects F-actin dependent cell migration (Fig. 8). Ori 3.1 cells were seeded into culture inserts to create a uniform wound of ∼500 μm, and wound closure was monitored for 9 hours. Whereas control cells closed ∼50% of the wound area after 9 h, IPZ-treated cells closed only 20% (Fig. 8A-C). In contrast, inhibiting Impα by IVM had no significant effect on wound closure, while combining IPZ and IVM had the same effect as IPZ alone (Fig. 8B). Consistent with reduced wound closure, IPZ- and IPZ/IVM-treated cells migrated significantly more slowly than control or IVM-treated cells (Fig. 8C). Impβ depletion by siRNAs (siKPNB1) similarly reduced wound closure efficiency and cell front velocity compared with control cells treated with non-targeting siRNAs (Fig. S8A and B). However, depletion of Impα (siKPNA2) or CRM1 (siXPO1), as well as inhibiting CRM1 nuclear export with KPT-330 or leptomycin B (LMB), had little to no effect on wound closure (Fig. S8A-D). Such behavior was also corroborated by quantifying in a separate experiment the migration velocity and distance covered by single cells with and without IPZ treatment (Fig. 8D and E; Supplemental Movies S2 and S3).

**Figure 8:**
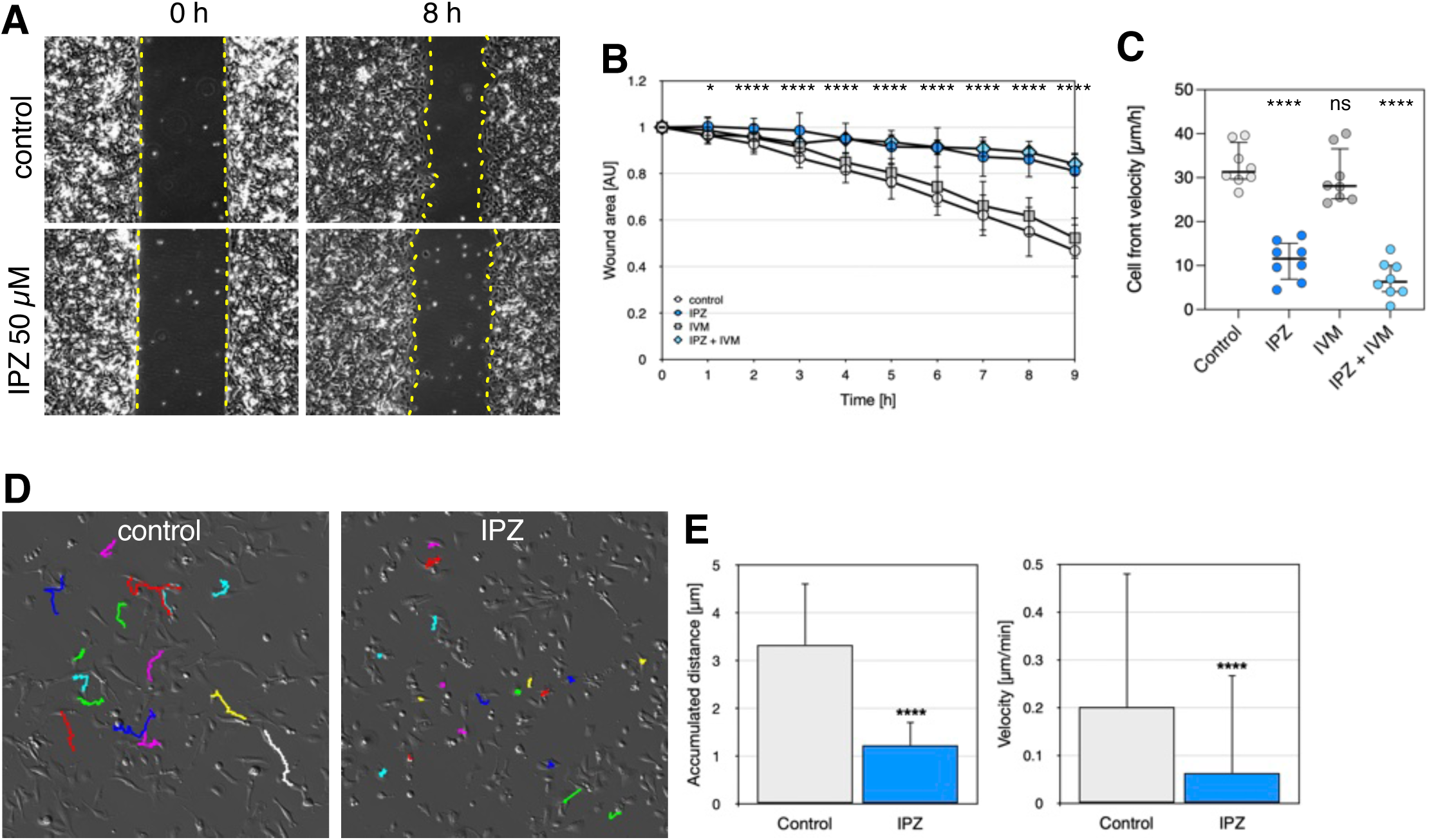
Inhibition of Impβ compromises collective and single cell migration. (**A**) Monitoring wound closure of Ori 3.1 cells (control) and cells treated with importazole (IPZ 50 µM, 1 h), Shown are representative examples at the indicated time points. Quantification of (**B**) the wound area and (**C**) the cell front velocity revealed that closure is significant reduced in Ori 3.1 cells treated with IPZ, as compared to control cells or cells treated with the importin-α inhibitor ivermectin (IVM). A combined treatment with IPZ and IVM affected collective cell migration of Ori 3.1 cells comparable to IPZ alone. (**D**) Live cell imagining of Ori 3.1 cells (control) and Ori 3.1 cells treated with IPZ (50 µM, 1 h) and trajectories of individual cell movement. Individual cells were manually tracked using Fiji/ImageJ. Quantification of (**E**) accumulated distance individual cells travel and the velocity. Inhibition of Impβ by IPZ affects cell movement and velocity significantly. Cells were imaged for 12 hours at 15 minutes intervals. Number of cells analyzed: n=40. Student’s T-test was used to calculate statistics: ****, p < 0.0001.

## Discussion

We have identified Impβ as a regulator of the actin cytoskeleton, separate from its established role in orchestrating NCT (Kalita et al., 2022; Kapinos et al., 2017; Kapinos et al., 2014; Wing et al., 2022). This is supported by the temporal separation of its effects: Impβ inhibition rapidly disrupted actin structures including stress fibers and filopodia, impairing cell migration within minutes, whereas nuclear import defects were detectable only after several hours. These kinetic differences indicate that the actin-related function of Impβ precedes NCT, consistent with rapid mechanosensitive actin remodeling versus slower, diffusion-driven NCT (Wang et al., 2009a).

Binding to actin may further act as a reservoir for Impβ whose engagement in NCT is coordinated with respect to changes in the cytoskeleton, resulting in the asymmetric partitioning of Impβ in the cytoplasm. Interestingly, loss of NCT partitioning as caused by abnormal NPC function or importin overexpression is associated with diseases with altered cellular mechanics including neurodegeneration (Coyne and Rothstein, 2022; Eftekharzadeh et al., 2018; Hall et al., 2021) and cancer (Agrawal et al., 2025; Alibert et al., 2017; Cagatay and Chook, 2018). Whether these associations reflect a causal relationship remains to be determined.

Our findings also resonate with long held views that Impβ perform functions beyond classical NCT (Harel and Forbes, 2004). The 19 HEAT repeats of Impβ coil into a short superhelix, thus providing extensive interaction surfaces on both the inside and the outside of the superhelix (Conti, 2002). Moreover, the intrinsic flexibility of Impβ may enable it to adapt to diverse binding motifs and molecules (Conti et al., 2006). Indeed, co-crystallization of Impβ with Impα, RanGTP, multiple FG-repeats, and various cargoes revealed that each partner engages distinct binding sites on Impβ (Bayliss et al., 2000; Cingolani et al., 2002; Cingolani et al., 1999; Lee et al., 2003; Vetter et al., 1999; Wohlwend et al., 2007). Regardless, the specific interface of Impβ:actin involves N-terminal residues of Impβ that are not engaged in nuclear import complexes (for review see: (Cingolani et al., 1999; Harel and Forbes, 2004)).

Actin is also known to bind a myriad of proteins. More than 400 actin-binding proteins (ABPs) have been identified thus far (Gao and Nakamura, 2022; Radulovic and Godovac-Zimmermann, 2011). Most ABPs have a typical binding motif, including the Wiskott-Aldrich syndrome (WASP)-homology domain-2 (WH2), the ADF/cofilin domain, or the calponin-homology (CH) domain, to interact with actin, while others use unique domains that are not based on sequence motifs (Dominguez, 2004; Gao and Nakamura, 2022). More generally, an actin-binding motif consists of an α-helix with few exposed hydrophobic side chains that binds in a hydrophobic cleft formed between actin subdomains 1 and 3 (Dominguez, 2004). This is exactly how Impβ may bind actin: based on our AF2 prediction, HEAT repeat 1 binds into the cleft between subdomain 1 and 3 of actin. Actin, while predominantly cytoplasmic, is also present in the nucleus (Dopie et al., 2012; Munsie et al., 2012; Ulferts et al., 2021; Vartiainen, 2008). As it lacks a classical NLS recognized by Impβ, the nuclear import of actin either requires cofilin (Dopie et al., 2012; Munsie et al., 2012) or through direct binding with Importin 9 (Keplinger et al., 2026). Thus, the formation of an Impβ:actin nuclear import complex is unlikely.

The interaction between Impβ and actin may have been overlooked in co-purification assays as actin’s interactions are often transient, dynamic and context specific (Viita et al., 2019). Particularly, binding affinities of regulatory proteins, such as the Arp2/3 complex, are in the weak, µM range, as shown by TIRF microscopy (Mullins et al., 1998). Myosin binding affinities for actin are context-dependent and range from 10 nm to 10 µM (Greene and Eisenberg, 1980a; Greene and Eisenberg, 1980b; Lymn and Taylor, 1971), depending on the nucleotide-binding state of the myosin head. However, proteomic studies have identified actin as Impβ interactor (Di Francesco et al., 2018; Song et al., 2022). Their association is evident in polarized cells, such as Ori 3.1 cells and fibroblasts, but less so in HeLa cells, possible due to their lack of a continuous cortical actin belt and reduced epithelial polarity (Braga et al., 1997; Ivanov et al., 2005; Zhang et al., 2005). Compared to conventional immunostaining, we find that smaller nano-secondary antibodies or primary antibodies directly conjugated with a fluorophore improve visualization and reveal clear co-localization of Impβ with actin stress fibers.

Interestingly, Impβ binds to G- and F-actin and its inhibition rapidly disrupts actin-based structures and functions, indicating a direct role in cytoskeletal organization. Consistent with this, Impβ inhibition by IPZ has been linked to altered actin dynamics and cell migration (Lu et al., 2024). These findings suggest that Impβ links cytoskeletal architecture with NCT, so as to balance fluctuating cytoskeletal forces with nuclear transport capacity. While “outside-in” mechanotransduction posits that mechanical inputs regulate nuclear import via changes in NPC diameter, our results introduce a complementary “inside out” mechanism in which the nuclear transport machinery itself modulates cytoskeletal organization. By binding actin and promoting filament assembly, Impβ may stabilize force-transmitting actin structures thereby tuning cytoskeleton tension and the mechanical environment of the nucleus. Together, this points to a reciprocal feedback loop in which cytoskeletal tension regulates nuclear transport, while transport receptors such as Impβ help maintain cytoskeletal organization and mechanical homeostasis. Further studies will be required to explore this behavior in more detail.

## Material and Methods

All experiments were carried out at room temperature unless otherwise stated. Analyses were performed in duplicate and were all repeated at least three times.

### Cell culture and transfection

Ori 3.1 cells were grown in RPMI-1640 (Merck Sigma-Aldrich, Buchs, Switzerland) supplemented with 10% fetal bovine serum (FBS) and 1% penicillin/streptomycin (pen/strep). MRC5, AG08468, and GM25336 fibroblasts were grown in Minimum Essential Medium (MEM) medium (ThermoFisher Scientific Gibco, Basel, Switzerland) supplemented with 15% fetal bovine serum (FBS) and 1% penicillin/streptomycin (pen/strep). HeLa, MDA-MB231, Rat2, and Rat2Sm9 cells were grown in Dulbecco’s modified Eagle medium (DMEM) supplemented with 10% FBS and 1% pen/strep. ARPE-19 cells were grown in DMEM-F12 medium supplemented with 5% FBS and 1% pen/strep. All cell lines were grown at 37°C in a humidified incubator with 5% CO_2_ atmosphere.

Plasmids were transfected using ViaFect (Promega, Dübendorf, Switzerland) and TurboFect transfection reagents (ThermoFisher Scientific) and siRNAs using Lipofectamine RNAiMax (ThermoFisher Scientific) following the instructions of the manufacturers. siRNAs were from Dharmacon (Lafayette, CO, USA): *KPNB1* (L-017523-00-0005), *KPNA2* (L-004702-00-0005), *XPO1* (L-003030-00-0005), and non-targeting siRNAs (D-001810-10).

### Drug treatment

For protein inhibition the following drugs were used: importazole (50 µM; SML0341-5MG, Sigma-Aldrich), ivermectin (2.5 µM; I8898-250MG, Sigma-Aldrich), leptomycin B (20 nM; L2915-5UG, Sigma-Aldrich), KTP-330 (Selinexor, 0.1 µM; S7252, Selleck Chemicals, Cologne, Germany), latrunculin A (0.2 µg/ml; 428026-50UG, Merck Millipore, Schaffhausen, Switzerland), cytochalasin-B (2 µg/ml; C2743-200UL, Sigma-Aldrich). For treatment of spheroids, importazole was used at 75 µM and ivermectin at 3.75 µM.

### Generation of stable cell lines

For the generation of Ori 3.1 cells expressing mClover-tagged endogenous importin-β, a CRISPR-based strategy exactly as described by (Yesbolatova et al., 2019) was used. In brief, to express the SpCas9 plasmid pX330-U6-Chimeric_BB-CBh-hSpCas9 (Addgene 42230; Supplemental Table S1) was used. Small guide RNA sequences to target exon 21 of *KPNB1* (the gene encoding importin-β) using primers 140/fw (Supplemental Table S2). To target *KPNB1* plasmids pMK289 (Addgene 72825; Supplemental Table S1) was modified: the auxin-inducible degron cassettes was first removed by In-fusion cloning. Next, a 1000 bp fragment of *KPNB1* was amplified from genomic DNA (HeLa cells) by PCR comprising 500 nucleotides upstream of the STOP codon and 500 nucleotides downstream of the STOP and cloned first into a conventional cloning vector, modified as described by Yesbolatova et al. and subcloned into the modified pMK289. Ori 3.1 cells were transfected with the two plasmids using Turbofect. 24 h after transfection, cells collected by trypsinization, resuspended in 1 ml growth medium and 100 µl cell suspension were plated in 10 cm dishes containing 10 ml medium. The next day, geneticin (700 µg/ml) was added to the medium. Medium was exchanged every 3-4 days. When visible, colonies were selected and cultured in a 96-well plate. Colonies were further cultured and genotyped by PCR. Clones harboring one *KPNB1* allele tagged were expanded and kept.

### Plasmids

Plasmids pDEST-ORF-V1 and pDEST-ORF-V2 (Addgene plasmids 73637 and 73638) were a gift of Darren Saunders (The Kinghorn Cancer Institute, Sydney, Australia), plasmid pX330-U6-Chimeric_NN-CBh-hSpCas9 (Addgene plasmid 42230) of Feng Zhang (Broad Institute of MIT and Harvard, Cambridge, MA, USA), plasmid pMK289 (Addgene 72827) of Masato Kanemaki (National Institute of Genetics, Mishimi, Japan), plasmids Dendra2-actin-C-18 (Addgene 57701), mCherry-vinculin-C-10 (Addgene 55014), mOrange-vinculin-23 (Addgene 57978) and mCherry-Arp2-N-14 (Addgene 54980) of Michael Davidson (Howard Hughes Medical Institute, Asburn, VA, USA), and pMRX-IPU-GFP-MYH9 (Addgene 168273) of Noboru Mizushima (University of Tokyo, Japan).

All constructs were generated by In-Fusion Cloning (Takara, Saint-Germain-en-Laye, France) using the manufacturer’s instructions. Mutations were inserted by site-directed mutagenesis using the QuikChange Lightning site-directed mutagenesis kit (Agilent Technologies, CA, USA) following the manufacturer’s instructions, deletions were inserted by In-fusion cloning. All constructs were verified by DNA sequencing. Plasmids used in this study are listed in Supplementary Table S1, primers are listed in Supplementary Table S2.

### Antibodies

The following antibodies were used as primary antibodies: mouse anti-Impβ (IF 1:500, WB 1:1000; clone 3E9, ab2811, Abcam, Cambridge, UK), mouse anti-actin (WB 1:2000; 66009-1-Ig, Proteintech), rabbit anti-actin (WB 1:1000; A-2066, Sigma-Aldrich), rabbit anti-YAP (IF: 1:100; ab52771, Abcam), rabbit anti-53BP1 (1:500; H-300, Santa Cruz Biotechnology, Heidelberg, Germany).

Secondary antibodies for immunofluorescence were: goat anti-mouse IgG-Alexa 488, goat anti-rabbit IgG-Alexa 488, goat anti-mouse IgG-Alexa 568, goat anti-rabbit IgG-Alexa 568 (Molecular Probes, Paisley, UK), goat anti-mouse abberior STARRed (STRED 1001, Abberior, Göttingen, Germany), and alpaca anti-mouse IgG2a CoraLite 488 (smg2a CL488-1-100, Proteintech). Phalloidin-iFluor 594 (1:1000; ab176757, Abcam) was used to stain F-actin. All goat antibodies were used at a dilution of 1:1000, except for STED antibodies, which were used at 1:200, all alpaca antibodies at 1:100. For CoraLite 488 labelling of the mouse anti-importin β antibody, the FlexAble CoraLite Plus 488 Labeling Kit for mouse IgG2a antibodies (KFA041, Proteintech) was used following the manufacturer’s instructions. For Western blot, secondary goat anti-mouse IgG and goat anti-rabbit coupled with alkaline phosphatase antibodies (Sigma-Aldrich) were used at a dilution of 1:10.000.

### Immunofluorescence microscopy

Cells were grown on glass coverslips and fixed with 2% formaldehyde for 15 min. Next, cells were washed three times with PBS for 5 min and permeabilized with PBS containing 0.2% Triton X-100 and 2% bovine serum albumin (BSA) for 10 min. After three washes with PBS containing 2% BSA for 5 min, cells were stained with the appropriate antibodies for 2 h at room temperature in a humid chamber. Next, cells were washed three times with PBS/2% BSA, incubated with secondary antibodies for 1 h and washed three times with PBS. The coverslips were then mounted with Mowiol-4088 (Sigma-Aldrich) containing 1 μg/ml DAPI and stored at 4°C until viewed. Images were acquired using a 63x oil immersion objective on a Leica TCS SP8 or a Leica Stellaris 8 Falcon point scanning confocal microscope (Leica Microsystems, Heerbrugg, Switzerland) or on an Axio Observer Z.7 microscope (Zeiss, Oberkochen, Germany). Images were recorded with the respective system software and processed using Fiji/Image J and Adobe Photoshop. All images were obtained close to the basal plane of cells except where specified.

### Permeabilized cell assays

Cells were grown on glass coverslips and washed twice with PBS for 5 minutes to remove growth medium. Next, cells were treated with 40 µg/ml digitonin (D141; Sigma-Aldrich) in ice-cold PBS for 5 min on ice. Cells were washed three times with PBS for 5 min, fixed in 2% formaldehyde for 15 min, washed three times in PBS containing 2% BSA for 5 min and stained and mounted as outlined above.

### Immunolabelling of spheroids

For the growth of spheroids, cells were collected by trypsinization, washed in fresh medium, and diluted to 5000 cells/ml. 100 µl cell suspension was plated per well of a 96-well ultra-low binding plate (Nunclon Sphera-treated, U-shaped bottom microplate; 174925, ThermoFisher Scientific) to make 500 cells per well. Spheroids were harvested on day 3 using a cut micropipette tip, transferred to a 48-well tissue culture plate, and fixed in 4% paraformaldehyde in PBS for 40 min at 4°C while shaking. Next, spheroids were permeabilized with PBS containing 0.5 % TritonX-100 and 3% BSA for 1 h while shaking, followed by incubation in blocking buffer (PBS containing 0.1 % TritonX-100, 3% BSA) for 2 h. 100 µl primary antibody solutions were added to each well and incubated overnight (∼12 h) with shaking. Adjacent free wells were filled with water to avoid evaporation of the antibody solution. Next, samples were washed 5 times 10 min with PBS on a shaking table, secondary antibodies were incubated over night at 4°C with shaking and samples were subsequently washed 5x 10 min in PBS. Glass slides were prepared for sample mounting using by adding 4 droplets of nail polish to prevent damage of the spheroids when mounting. After washing, 20 µl of Vectashield mounting medium (Vector Laboratories, Lubio Science, Zurcih, Switzerland) was added to each spheroid, spheroids in Vectashield were transferred to the center between the drops of nail polish drops, covered with a glass coverslip, and sealed with nail polish. Images were recorded on an Olympus SpinD or SpinSR spinning disc microscope (Olympus Switzerland AG, Wallisellen, Switzerland) using the system software and processed using Fiji/Image J and Adobe Photoshop.

### Bimolecular fluorescence complementation assays

Ori 3.1 and HeLa cells, respectively, were grown in 24-well plates on cover slips and transfected with 250 ng of each plasmid using ViaFect (Ori 3.1) or Turbofect (HeLa) transfection reagent. After 24 hours, cells were fixed in 2% formaldehyde in PBS for 15 min, washed in PBS, permeabilized with 0.2% Triton X-100 in PBS containing 2% BSA for 10 min. Blocking was performed with 2% BSA in PBS for 30 min and cells were incubated with phalloidin-iFluor 594 reagent for 1 hour. After three washes in PBS, samples were mounted on glass slides using Mowiol containing 1 µg/ml DAPI and stored at 4°C until viewed. Cells were imaged using a confocal point scanning microscope (Leica TCS SP8, Leica Microsystems, Heerbrugg, Switzerland). Images were recorded using the microscope system software with uniform laser settings for all samples. Fluorescence intensity was measured using the Fiji/Image J. Images were processed using Fiji/Image J and Adobe Photoshop.

### Expression and purification of Importin-β, fascin, and mDia2

All recombinant proteins were produced in *E. coli* BL21 codon plus (DE3) cells. Bacteria precultures were grown overnight at 37°C in 100 ml LB medium containing appropriate antibiotics, 10 ml of the preculture were diluted into 1 l LB medium (the total culture volume was 3 l) and growth temperature was reduced to 21°C. Protein expression was induced with 0.5 mM of isopropyl-beta-D-thiogalactopyranoside (IPTG) at an optical density at 600 nm of 0.5 and cells were grown over night at 21°C. The cells were collected by centrifugation at 8°C at 5.422 g for 20 min. The pellet can be frozen at -20°C or directly processed for protein purification.

Next the pellets were resuspended in 10 ml of the appropriate lysis buffer (Impβ and fascin: 10 mM Tris, pH 7.5, 100 mM NaCl, 1 mM DTT, 10 mM imidazole; mDia2: 50 mM Tris, pH 8.0, 300 mM NaCl, 1 mM EDTA, 1 mM DTT, 10 mM imidazole). All lysis buffers were supplemented with 1.5 µg/ml DNaseI (10104159001; Roche, Basel, Switzerland), 1 mg/ml lysozyme (A4972; AppliChem, Darmstadt, Germany), 1 mg/ml Pefabloc (76307; Sigma-Aldrich), and cOmplete proteinase inhibitor cocktail (11 836 145 001; Roche). The resuspended samples were incubated for 1 h at 4°C on a roller mixer and subsequently sonicated on ice (30 % output, over 5 min with 30 sec on and 4 sec off intervals; Branson 550 Sonifier, Shanghai, China). After centrifugation at 4°C at 25.000 g for 1 h, the supernatants were collected, loaded onto a 5 ml HisTrap HP Cytiva column (GE17-5248; Sigma-Aldrich), and proteins were eluted with increasing concentrations of imidazole (10-500 mM). Collected fractions were dialyzed for 1 h at 4°C against 4 l buffer (10 mM HEPES, pH 7.2, 200 mM NaCl). The dialyzed fractions were transferred into a 50-ml Falcon tube, concentrated to 3 ml, and loaded onto a size exclusion column (HiLoad 16/600 Superdex 75 pg; Cytiva 28-9893-33; Sigma-Aldrich) using an ÄKTA protein purification system (Cytiva; Sigma-Aldrich). Proteins were eluted using with 10 mM Tris, pH 7.5, 100 mM NaCl, 1 mM DTT (importin-β); PBS (fascin); 50 mM Tris, pH 8.0, 300 mM NaCl, 1 mM EDTA 1 mM DTT (mDia2). Collected fractions were concentrated to appropriate concentrations, frozen in liquid nitrogen, and stored at -80°C.

### Expression and purification of human β-actin

Human β-actin was expressed using a cold shock vector according to (Tamura et al., 2011). *E. coli* BL21 codon plus (DE3) precultures transformed with pCold1-actin were grown as outlined above. When OD_600_ of 0.5 was reached, cultures were rapidly cooled down by placing the Erlenmeyer flask on ice for 1 h. Next the Erlenmeyer flask was returned to the shaking incubator set to 15°C, grown for another 30 min before protein expression was induced by 50 µM IPTG. Cultures were grown over night at 15°C. The cells were collected by centrifugation at 8°C at 5422 g for 20 min and the pellet frozen at -20°C.

For actin purification, the pellet was resuspended in buffer A (50 mM Tris, pH 8.0, 1 mM EDTA, 1 mM DTT, 1 mM ADP) supplemented with DNAseI, lysozyme, Pefabloc and protease inhibitor cocktail, as above. Samples were incubated for 1 h at 4°C while rolling, sonicated (30%, 5min, 3sec on, 4sec off), and collected by centrifugation at 30.000 g, for 30 min at 4°C. The supernatant was transferred into a 50-ml Falcon tube, 6 ml of a 50% Ni-NTA bead slurry (635660, Takara) were added, and incubated over night at 4°C while rolling.

Next, samples were centrifuged at 500 g for 10 min at 4°C and the pellet resuspended in buffer B (50 mM Tris, pH 8.0, 1 mM EDTA, 1 mM ADP (freshly added)), spun again at 500 g for 10 min at 4°C, the pellet was resuspended into 20 ml 0.5 % (v/v) Triton-X100 in sterile water supplemented with 1 mM ADP. Samples were spun as before and the pellet next resuspended in buffer B containing 10 mM imidazole, and spun again. Next, the pellet was resuspended in 10 ml buffer B containing 1 mM DTT and 250 mM imidazole, samples were spun again, the supernatant was collected. These last 3 steps were repeated twice, so that a total volume of about 30 ml supernatant was collected and 50 µl of Pefabloc were added. The supernatant was dialyzed for 2 h at 4°C against 2 l buffer G (5 mM Tris, pH 8.0, 0.2 mM CaCl_2_, 0.2 mM ATP, 0.1 mM DTT). Dialysis was repeated a second time into fresh 2 l of buffer G and samples concentrated to 2 ml using a 15 ml sample volume Amicon Ultra, 10 kDa centrifugal filter (UFC9010, Merck-Millipore). Next, a 5-ml Ni-column (GE17-5248; Sigma-Aldrich) was equilibrated with buffer G containing 20 mM imidazole, the concentrated sample solution loaded and incubated for 1 h at 4°C. The column was washed with 20 ml buffer G (1 ml/min) and actin was purified on a size exclusion column as outlined above, using a gradient of imidazole (20-100 mM) in buffer G. Collected fractions were dialyzed over night against 3 l of buffer G at 4°C. After dialysis, the collected fractions were concentrated to 1 ml, and added on top to a 1.5 ml ultracentrifugation tube (343778, Beckman Coulter) containing 300 µl of buffer DG (5 mM Tris, pH 8.0, 0.2 mM CaCl_2_, 0.1 mM DTT, 10% glycerol), and centrifuged for 1 h at 265.000 g at 24°C. Finally, the supernatant was collected, stored on ice, and lyophilized with vacuum at 45°C for 12 h. For long term storage, actin was reconstituted to 10 mg/ml in deionized water, aliquoted to 5 µl, snap frozen in liquid nitrogen, and stored at -80°C.

### Labelling of Impβ for STED imaging

To label recombinant Impβ for STED imaging, a three-fold molar excess of Abberior STAR 635P NHS-ester dye (ST635P-0002; Abberior, Göttingen, Germany) was added and incubated for 1 h in the dark, with gentle mixing. Next, the mixture was transferred to a spin column (Centri Pure Mini XDesalt Z-50; CP-0219, emp Biotech, Berlin Germany) and spun at 2.400 g for 5 min to remove unbound dye. The retentate containing labeled Impβ was collected and protein concentration as well as dye incorporation were determined using a NanoDrop One spectrometer (ThermoFisher Scientific). The absorbance values at 280 nm and the dye-specific wavelength (λ = 635 nm) were used to calculate protein concentration and the degree of labeling (DOL).

### GST-pull down assays

All recombinant GST and GST-Impβ were produced in *E. coli* BL21 codon plus (DE3) cells. Bacteria pre-cultures were grown overnight at 37°C in 1 ml LB medium containing appropriate antibiotics and diluted into 100 ml LB medium containing appropriate antibiotics. Protein expression was induced with 0.1 mM of IPTG at an optical density at 600 nm of 0.5 and cells were grown for further 3 h at 37°C. The cells were collected by centrifugation at 4°C at 3.220 g for 20 min, resuspended in 4 ml PBS containing 1% Triton X-100 plus protease inhibitor cocktail and next sonicated on ice (5 times 10 sec, with 10 sec off between each sonication). After centrifugation at 4°C at 16.000 g for 15 min, the supernatants were collected, frozen in liquid nitrogen, and stored at -80°C.

For the preparation of cell lysates for pull-down assays, Ori 3.1 cells were grown to confluency in a T75 tissue culture flask. Cells were harvested by scrapping, washed in PBS, resuspended in 400 µl lysis buffer (50 mM Tris-HCl, pH 7.4, 250 mM NaCl, 0.1 % Triton X-100, 2 mM EDTA-Na_2_, 10% glycerol, protease inhibitor tablets), and frozen at -20°C.

For pull-down assays, 100 μl of GST (plus 150 µl 2x binding buffer (2xPBS, containing 10% glycerol and 1% Triton X-100)) or 250 µl GST-fusion proteins were bound to 25 μl of magnetic glutathione beads for 1 h at 4°C on a rocker platform. Next, the beads were washed twice with 2x binding buffer and subsequently incubated with 100 µl Ori 3.1 lysate plus 150 µl 2x binding buffer overnight at 4°C on a rocker platform. After binding, 50 µl of the unbound fractions were collected, and beads were washed twice with 2x binding buffer and followed by two washes with detergent-free 2x binding buffer. To elute bound proteins, beads were resuspended in 60 µl 2x Laemmli buffer (125 mM Tris-HCl, pH 6.8, 4% SDS, 20% glycerol, 10% 2-mercaptoethanol, 0.004% bromophenol blue) and boiled for 5 min at 95°C. Next, the samples were separated by SDS-PAGE and analyzed by Western blot. 20 µl of each, the unbound and bound fractions were loaded.

### In vitro binding assays

For *in vitro* binding assays, actin was first depolymerized by resuspending the stock solution to 2.5 mg/ml in General Actin buffer (5 mM Tris-HCl, pH 8.0, 0.2 mM CaCl_2_) supplemented with 0.2 mM ATP and 0.5 mM DTT. After incubation for 60 min on ice, the protein was centrifuged for 15 min in a microcentrifuge at top speed. The supernatant was used for the assays.

GST fusion proteins were expressed and processed as outline above. After immobilization of the GST fusion proteins on the beads and washing, the beads were next incubated overnight with the respective recombinant proteins (25 µg in 250 µl 2x binding buffer) at 4°C on a rocker platform. After binding, 50 µl of the non-bound fractions were collected, beads washed as described above, bound fractions eluted, and samples separated by SDS-PAGE and analyzed by Western blot as described above. For actin binding assays, 250 µl General Actin binding buffer was used instead of 2x binding buffer, and samples were washed twice with 2x binding buffer, two times with detergent-free 2x binding buffer, and once with 500 mM NaCl prior to elution with sample buffer.

### Actin polymerization assays

Lyophilized rabbit skeletal muscle actin (A2522; Sigma-Aldrich) was reconstituted in deionized water to a concentration of 10 mg/ml, snap frozen in liquid nitrogen and stored at - 70°C. For co-sedimentation assays, actin was resuspended and diluted in general actin buffer (5 mM Tris-HCl, pH 8.0, 0.2 mM CaCl_2_) supplemented with 0.2 mM ATP and 0.5 mM DTT to a concentration of 1 mg/ml (21 µM). The resuspended actin was incubated on ice for 60 min to depolymerize actin oligomers formed during storage (G-actin). For the preparation of F-actin, the actin was next mixed 1:10 with 10x actin polymerization buffer (500 mM KCl, 20 mM MgCl_2_, 10 mM ATP) and incubated for 1 h at room temperature. Test proteins (Impβ, mDia2, fascin) were centrifuged at 150.000 g for 1 h at 4°C to remove potential aggregates formed during storage. To test for G-actin binding, Impβ and mDia2, respectively, were incubated with G-actin for 30 min at RT. The final concentrations were 16.8 µM for actin and 6 µM for Impβ and mDia2. Actin polymerization was next induced by adding a reduced amount of actin polymerization buffer resulting in 0.25x concentration. Samples were incubated exactly 30 min. To test for F-actin binding, Impβ and fascin, respectively, were incubated with F-actin for 30 min. The final protein concentrations in the assays were 16.8 µM for actin, 6 µM for Impβ and mDia2, and 4 µM for fascin. After incubation, samples were centrifuged at 150.000 g for 1.5 h at 24°C. Equal amounts of the supernatant and the pelleted fractions were next analyzed by 12% SDS-PAGE and Coomassie-Blue staining.

For kinetic measurements, actin polymerization was monitored by the pyrene assay in which pyrene-labeled actin was added to 5% of the total actin concentration (Qu et al., 2015). In brief, actin (5% labelled) was depolymerized for 1h as above and next centrifuged for 30 min at 150.000 g at 4°C. The supernatant was taken and 10% of 10x magnesium exchange buffer (500 μM MgCl_2_, 2 μM EGTA) was added 30 min prior to measurement. 30 μl of the Mg-actin (5% labelled) were pipetted into six wells of a black polystyrene 96-well plate. In another six wells 88.4 μl of G-buffer were mixed with 10 μl of 10x KMEI buffer (500 mM KCl, 100 mM imidazole, pH 7.0, 10 mM MgCl_2_, 10 mM EGTA, pH 7.0) and 1.6 μl of either Impβ or of Impβ buffer (10 mM Tris, pH 7.5, 100mM NaCl, 1mM DTT, 10mM Imidazole). 70 μl of the polymerization mixture was added to the wells containing 30 μl of Mg-actin (5% labelled) to yield a final concentration of 0.5 μM Mg-actin (5% labelled), 1μM Impβ and 0.7x KMEI-buffer. After mixing using a multichannel-pipette, the plate was immediately transferred to a Tecan Spark Multi-Mode Plate-Reader (Tecan Austria GmbH, Grödig, Austria) and the fluorescence increase of pyrene-actin was monitored for 2000 sec using a Tecan Spark Multi-Mode plate reader (Tecan Austria GmbH, Grödig, Austria) at wavelength settings for excitation and emission of 365 nm and 407 nm, respectively.

For imaging, phalloidin-iFluor594 (1:1000) was added to the pellet fractions and incubated for 10 min at RT. For confocal imaging, samples were placed on glass sides and mounted with Mowiol, and for STED imaging with Prolong Diamond Antifade mounting medium (Invitrogen). For negative staining EM, 5 µl of each sample were placed on freshly glow-discharged copper grid (mesh 200) covered with parlodium/C film for 1 min and washed three times with water and once with 2% uranyl acetate. For negative staining, the samples were incubated for 30 s with 2% uranyl acetate. Excess staining solution was removed with filter paper and, subsequently, the grids were air dried. Three different grids were prepared for each sample. The grids were analyzed using a transmission electron microscope (Philipps CM-100) operated at an acceleration voltage of 80 kV.

### STED imaging

STED images were acquired on a Leica Stellaris 8 Falcon STED microscope equipped with a pulsed 775 nm depletion laser and a pulsed white light laser (WLL) for excitation. For acquiring images, a 100x oil immersion objective lens (NA 1.4) was used. Images were acquired and processed using the Leica LASX software. Images were taken in frame mode with an optical zoom of 5, image size was 23.25 μm with a pixel size of 19.92 nm.

### Western blots

Cells were harvested by trypsinization, washed with PBS, and resuspended in lysis buffer (50 mM Tris-HCl, pH 7.5, 150 mM NaCl, 1% NP-40) supplemented with a cocktail of protease inhibitor tablets (Roche) and frozen at -20°C. Subsequently, the lysates were cleared by centrifugation at 16.000 g at 4°C for 5 min. The lysates were resuspended in Laemmli buffer and boiled for 5 min at 95°C. 30 µg of proteins were then separated by SDS-PAGE and subsequently transferred onto a PVDF membrane with a current of 100 mA for 1 h. Next, the membrane was incubated for 1 h in PBS containing 0.1% Tween 20 (PBS-T) and 0.1% of Tropix I-Block (Applied Biosystems, T2015), followed by incubation of primary antibodies in the blocking solution for overnight at 4°C. After washing three times with PBS-T, the membrane was incubated with the appropriate alkaline phosphatase-conjugated secondary antibody for 1 h. After three washes with PBS-T, the membrane was washed twice with assay buffer (100 mM Tris-HCl, pH 9.8, 10 mM MgCl_2_) for 2 min. The membrane was incubated for 5 min with the Lightning CDP Star Chemiluminescence reagent (ThermoFisher Scientific, Applied Biosystem) and visualized using a Fusion FX chemiluminescence imager (Vilber, Marne-la-Vallée, France).

### Wound-healing assays and single cell tracking

Ori 3.1 cells were seeded into Culture-Insert 2 Well (Ibidi, Gräfelfing, Germany) and grown overnight. For drug treatment, the respective inhibitor was added to the growth medium at the above indicated concentrations and incubated for 1 h prior to imaging. For siRNA treatments, siRNAs were transfected as described above and incubated for 48 h. Inserts were removed and wound closure was monitored every hour using a Leica PAULA microscope. Wound area and cell front velocity were analyzed using Fiji/ImageJ.

For single cell tracking, cells were plated into 6 cm culture dishes and grown overnight to about 30% confluency. The medium was replaced by phenol red-free medium, the cells incubated for at least 3 h in the phenol red-free medium. Drugs were then added as described above and imaging started immediately. Movies were recorded for 12 h at 15 min intervals on a Leica spinning-disc confocal or a Leica MICA wide-field microscope. For analysis, individual cells were tracked manually using the Fiji/Image J plug in Cell tracking and the accumulated distance individual cells travel and their velocity were calculated.

### Surface plasmon resonance measurements

SPR experiments were performed at 25°C on a Biacore T200 instrument (Cytiva, Glattbrugg, Switzerland), as previously described (Kapinos et al., 2017; Kapinos et al., 2014; Schoch et al., 2012). In brief, Impβ and the N145 fragment of Impβ were immobilized on a CM5 sensor chip (BR100012, Cytiva) to a surface density of ∼300–1200 RU using standard amine coupling chemistry. Importazole (IPZ) was serially diluted (1:1) in running buffer (1x PBS, 1mM MgCl2, 1mM TCEP) containing 5% DMSO and injected at concentrations of 0-100 µM for Impβ and 0-50 µM for the N145 fragment. Equilibrium responses (R_eq_) were plotted as a function of increasing concentrations and fitted to a one-component binding Langmuir isotherm using Biacore Evaluation software (individual fits for each replicate). Four independent experiments were performed.

### Statistics

Plots and statistics were established using GraphPad Prism 10 (GraphPad Software, Boston, MA, USA) or Apple Numbers. Statistics were calculated on averaged values of biological replicates, according to (Lord et al., 2020). Two-Way Anova tests were carried out to determine *p*-values.

## Acknowledgements

The authors thank Dr. Elsa Obergfell (Biozentrum, University of Basel, Switzerland) for help with Alphafold2 modelling. We thank Drs. Darren Saunders (The Kinghorn Cancer Institute, Sydney, Australia), Feng Zhang (Broad Institute of MIT and Harvard, Cambridge, MA, USA), Michael Davidson (Howard Hughes Medical Institute, Asburn, VA, USA), Channing Der (Lineberger Comprehensive Cancer Center, Chapel Hill, NC, USA), Joel Swanson (University of Michigan Medical School, Ann Arbor, MI, USA), Masato Kanemaki (National Institute of Genetics, Mishimi, Japan), and Noboru Mizushima (University of Tokyo, Japan) for sharing reagents. Dr. Ramona Jühlen (RWTH Aachen, Germany) is acknowledged for suggestions and discussions, and Antonia van den Broek, Elias Engelhardt and Anna Gross for technical assistance. Confocal images were acquired at the Imaging Core Facility, Biozentrum, University of Basel.

This research was supported by the Biozentrum and the Swiss Nanoscience Institute. R.Y.H.L. acknowledges funding support from the Swiss National Science Foundation (Swiss National Science Foundation/COST (grant IZCOZ0_198146).

