## Supplemental Material for "The nuclear transport receptor Impβ is a regulator of actin polymerization"

### Supplementary information

#### 1. Supplementary methods

##### *Live cell imaging and single cell tracking*

For single cell tracking, cells were plated into 6 cm culture dishes and grown overnight to about 30% confluency. The medium was replaced by phenol red-free medium, the cells incubated for at least 3 h in the phenol red-free medium. Drugs were then added as described above and imaging started immediately. Movies were recorded for 12 h at 15 min intervals on a Leica spinning-disc confocal or a Leica MICA wide-field microscope. For analysis, individual cells were tracked manually using the Fiji/Image J plugin "*Cell tracking*" and the accumulated distance individual cells traveled and their velocity were calculated.

#### 2. Figure legends

**Figure S1**: ***Localization of Impβ in distinct human cell lines.*** (**A**) MRC5 fibroblasts were immunostained with for Impβ using secondary Alexa 488 conjugated antibodies (Secondary A488; left), Nano-secondary Alexa 488 conjugated secondary antibodies (Nano-secondary; middle), or CoraLite488 conjugated primary antibody (Primary CoraLite 488; right). Impβ localizes to the nucleus, the cytoplasm, and nuclear rim, and enriches at the cell cortex and associates with actin stress-fibers becomes. (**B**) Impβ is enriched at the leading edge of the fibroblast cell line AG08648, in ARPE-19 primary epithelial cells as well as in MDA-MB231 triple-negative breast cancer cells and in HeLa cervix cancer cells. The association of Impβ with the cell cortex does not depend on the support or the presence of actin stress fibers: it can be detected at the cell cortex in cells grown on collagen IV (col IV) coated cover slips (**C**) and upon actin depolymerization (**D**) by Y27632, cytochalasin B (cytoB) or latrunculin A (latA). Shown are representative confocal images. Scale bars, 10 µm.

**Figure S2**: ***Control experiments for bimolecular fluorescence complementation assays.*** BiFC signal produced by interaction between (**A**) Impβ and actin in HeLa cells. Actin interacts with full-length Impβ (Impβ:actin), the N-terminal 145 residues (N145:actin), and the N-terminal 31 residues (N31:actin). Deletion of residues 2-31 (i.e., HEAT repeat 1; N145Δ2-31:actin) largely abolished the interaction of Impβ with actin. Negative controls for the BiFC assays in Ori 3.1 cells: (**B**) isoform 2 of Impβ (Iso2:actin), Impβ and the actin-related protein Arp2 (Impβ:Arp2), as well as Impβ and tubulin (Impβ:tubulin); (**C**) Impα:actin, Impβ2:actin, XPO1:actin, Ran:actin. Positive controls included (**D**) Impβ:Impα, actin:vinculin, and actin:myosin. (**E**) Localization of Impβ-EGFP fusion proteins in Ori 3.1

and HeLa cells. Shown are representative confocal images. DNA was visualized by DAPI. Scale bars, 10 μm.

**Figure S3**: ***Control experiments for bimolecular fluorescence complementation assays.*** BiFC signal produced by interaction between Impβ N31 and actin as well as mutants thereof in (**A**) Ori 3.1 cells and (**B**) HeLa cells. Actin interacts with the N-terminal 31 (N31:actin) residues of Impβ. Mutating E8 and Q22 of Impβ to alanine (N31-E8AQ22A) and S350 and E167 of actin (actin-E167AS350A) largely reduced the interaction of Impβ with actin, while mutations in all four residues (N31-E8AQ22A:actin-E167AS350A) abolished the interaction. (**C**) BiFC signal originated from an interaction between full length Impβ and actin in Ori 3.1 cells. Mutating E8 and Q22 of Impβ to alanine (Impβ-E8AQ22A) and S350 and E167 of actin (actin-E167AS350A) largely reduced the interaction of Impβ with actin, while mutations in all four residues (Impβ-E8AQ22A:actin-E167AS350A) abolished the interaction. DNA was stained with DAPI (blue). Shown are representative confocal images. Scale bars, 10 μm.

**Figure S4:** ***Importazole binds to the N-terminal region of Impβ and comprises its binding to actin.*** (**A**) AlphaFold3 modelling of the Impβ-IPZ interface. (**B**) Representative Surface plasmon resonance (SPR) sensograms revealing the binding of IPZ to Impβ (left) as well as to the Impβ N145 (middle). Average equilibrium binding constants (KDs) for Impβ (black) and Impβ N145 (blue) were similarly about 40 μM, as determined by fitting to the Langmuir binding isotherm (left). Shown are non-fit response curves representing 4 replicates. Mean values are indicated by circle symbols. Error bars represent ±SD. (**C**) SDS-PAGE analysis of the effect the respective pre-incubation of Impβ as well as actin with IPZ. Impβ was incubated for 30 minutes with 100 μM IPZ prior to incubation with F-actin. F-actin alone was treated alike. The supernatant (S) and pellet (P) fractions after high-speed centrifugation were analyzed by SDS-PAGE and (**D**) the respective levels of actin in pelleted fractions were quantified. Number of assays: n=4.

***Figure S5**: **Inhibition/depletion of Impβ affects actin stress fibers***. Stress fiber detected by the FilamentSensor II corresponding to the confocal images (**A**) shown in Figure 4A. (**B**) Ori 3.1 expressing Impβ-mClover cells were grown on glass cover-slips and treated with 50 μM importazole (IPZ) for respectively 5 min and 60 min. Cells were stained with phalloidin to visualize F-actin. Shown are representative confocal images. Actin stress fibers and Impβ-mClover imprints were revealed using the open-source JRE analysis tool FilamentSensor II. Quantitative analysis of actin stress fibers summarizing (**C**) the number of actin filaments per cell, as well as (**D**) the length and (**E**) the width of actin filaments. Quantitative analysis of Impβ-mClover imprints on actin summarizing (**F**) the number of actin filaments per cell, as well as (**G**) the length and (**H**) the width of the filaments.

**Figure S6:** ***Importazole treatment affects nuclear import and Impβ distribution only after several hours.*** Ori 3.1 cells were grown on glass cover-slips and respectively treated with 50 μM importazole (IPZ) for respectively 5 min (IPZ5), 15 min (IPZ15), 30 min (IPZ30), 60 min (IPZ60), and 360 min (IPZ360). (**A**) Nuclear import of the Impα/Impβ cargo 53BP1 was monitored by confocal microscopy, as well as the localization of (**B**) Impβ itself. Their respective nuclear-to-cytoplasmic ratio as determined using Fiji/ImageJ (**C-D**). The number of analyzed cells is indicated at the top of each column. Two-Way Anova test was used to calculate statistics: ***, $p < 0.001$; ns, non-significant.

**Figure S7**: ***Inhibition of Impβ compromises cell migration.*** Quantification of (**A**) wound closure and (**B**) cell front velocity in Ori 3.1 cells treated with siRNAs targeting Impβ (siKPNB1), Impα (siKPNA2), or the exportin CRM1/XPO1 (siXPO1): depletion of Impα or CRM1 had no or little effect on collective cell migration of Ori 3.1 cells, in contrast to depletion of Impβ. siNT, non-targeting control siRNAs. Number of assays: N=6-12. Scale bar, 100 µm. (**C**) Inhibition of CRM1 has no obvious effect on collective migration of Ori 3.1 cells in wound healing assays. Quantification of the wound area and (**D**) the cell front velocity in Ori 3.1 cells treated with KPT-330 or LMB. Two-Way Anova test was used to calculate statistics: *, $p < 0.05$, **, $p < 0.01$, ***, $p < 0.001$, ****, $p < 0.0001$; ns, non-significant.

**3. Supplementary movies**

***Movie 1***: Ori 3.1 Impβ-mClover. Movie was recorded on a Zeiss LSM980 laser scanning confocal microscope equipped with AiryScan2 detector. Movie was recorded for 8 h at 15 min intervals.

***Movie 2***: Live cell imaging of Ori 3.1 for single cell tracking. Movie was recorded on a Lecia Mica microscope for 12 h at 15 min intervals

***Movie 3***: Live cell imaging of Ori 3.1 treated with 50 µM importazole for single cell tracking. Movie was recorded on a Lecia Mica microscope for 12 h at 15 min intervals

### Figure S1

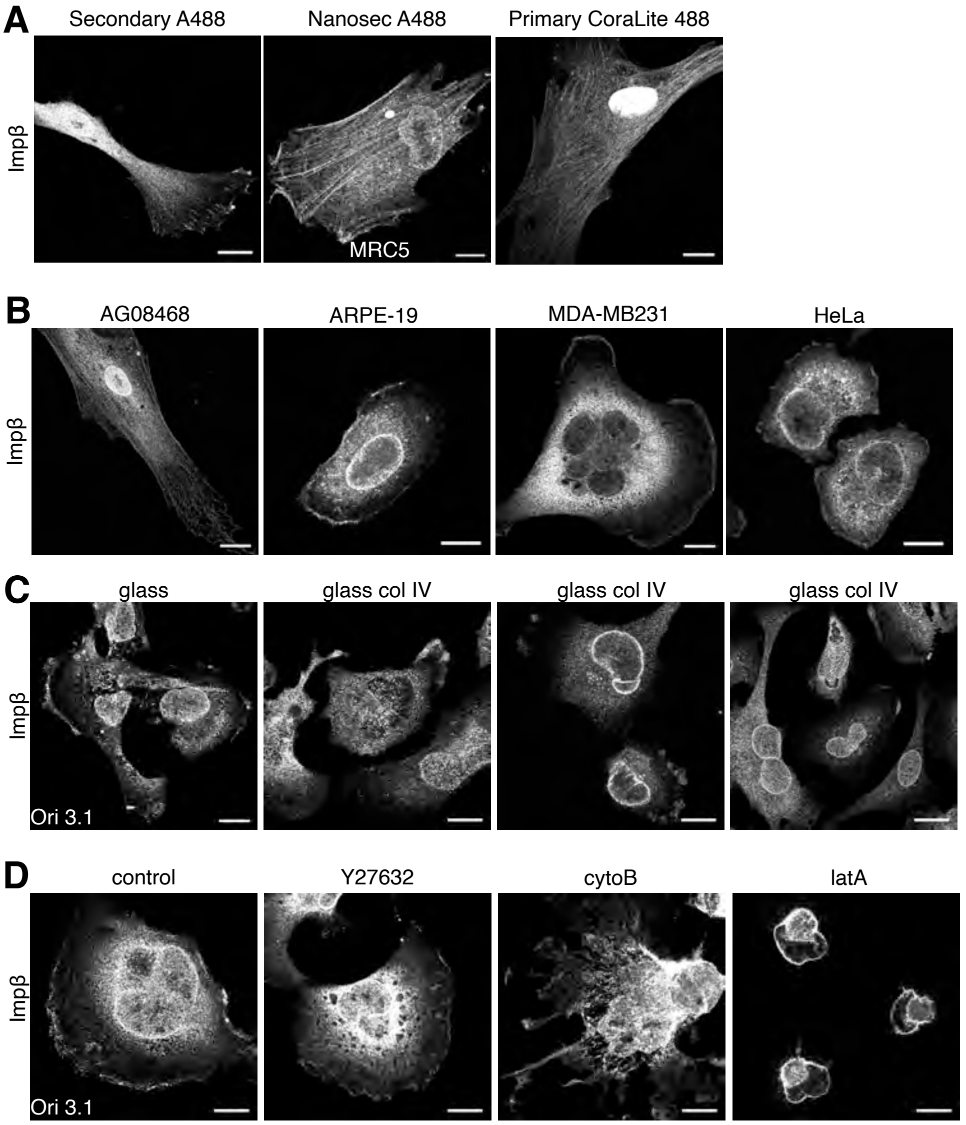
A
Secondary A488
Nanosec A488
Primary CoraLite 488
Impβ
MRC5
B
AG08468
ARPE-19
MDA-MB231
HeLa
Impβ
C
glass
glass col IV
glass col IV
glass col IV
Impβ
Ori 3.1
D
control
Y27632
cytoB
latA
Impβ
Ori 3.1

### Figure S2

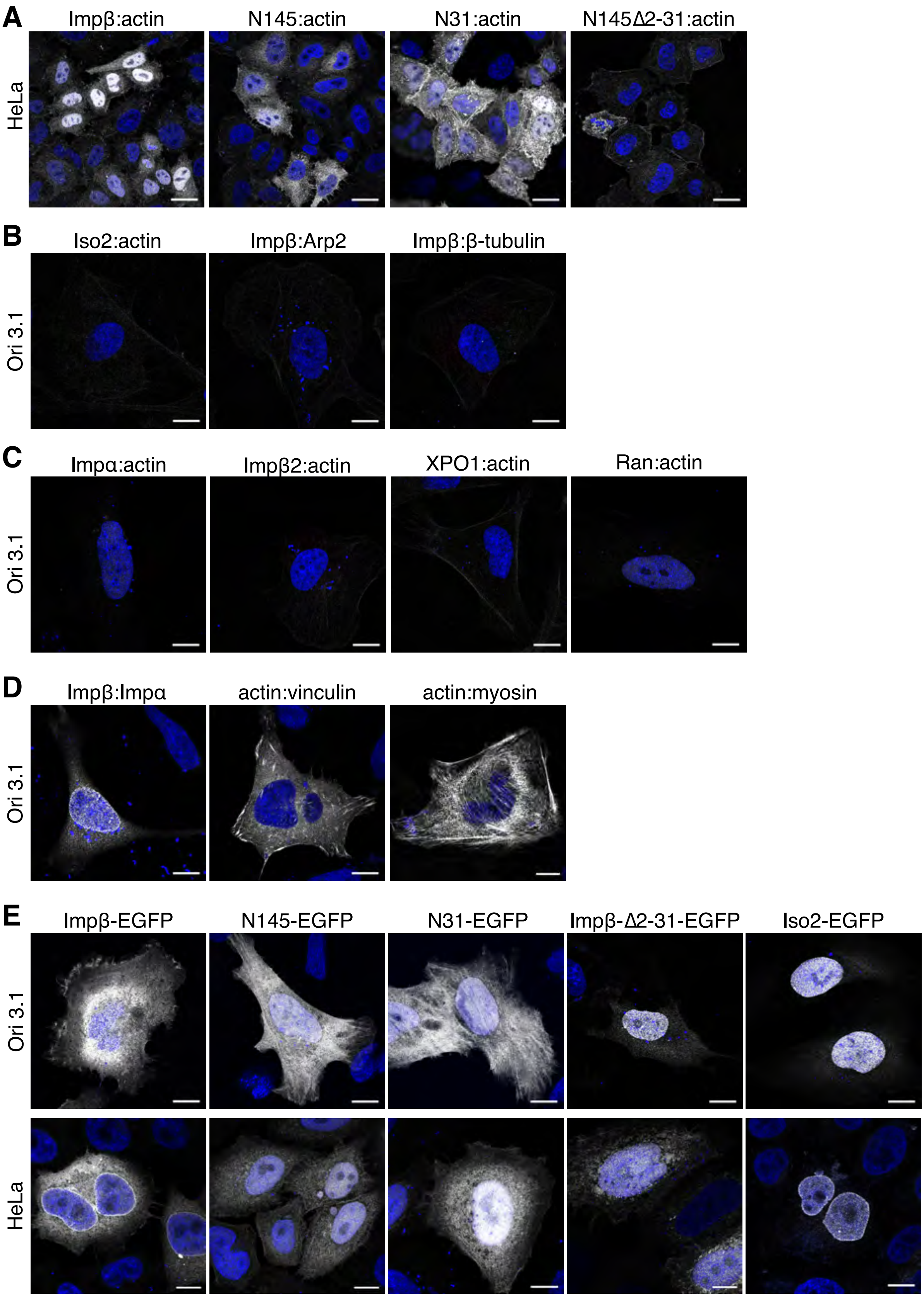
A
Impβ:actin
N145:actin
N31:actin
N145Δ2-31:actin
HeLa
B
Iso2:actin
Impβ:Arp2
Impβ:β-tubulin
Ori 3.1
C
Impα:actin
Impβ2:actin
XPO1:actin
Ran:actin
Ori 3.1
D
Impβ:Impα
actin:vinculin
actin:myosin
Ori 3.1
E
Impβ-EGFP
N145-EGFP
N31-EGFP
Impβ-Δ2-31-EGFP
Iso2-EGFP
Ori 3.1
HeLa

### Figure S3

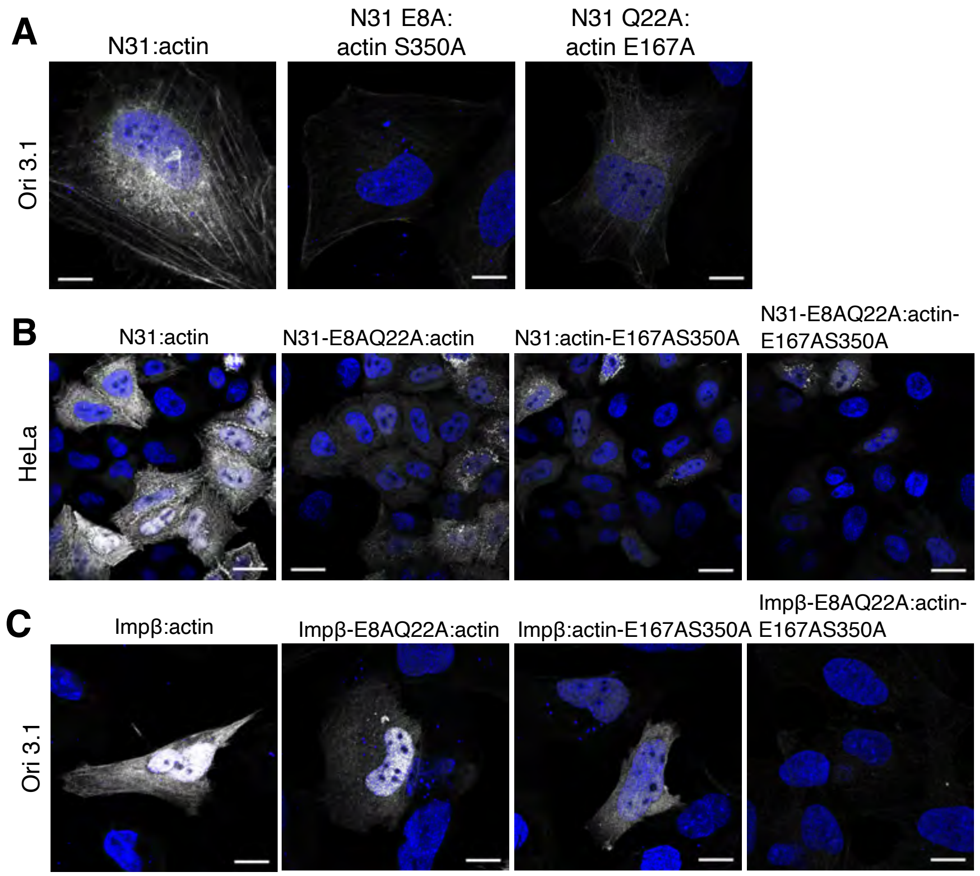
A
N31:actin
N31 E8A:
actin S350A
N31 Q22A:
actin E167A
Ori 3.1
B
N31:actin
N31-E8AQ22A:actin
N31:actin-E167AS350A
N31-E8AQ22A:actin-
E167AS350A
HeLa
C
Impβ:actin
Impβ-E8AQ22A:actin
Impβ:actin-E167AS350A
Impβ-E8AQ22A:actin-
E167AS350A
Ori 3.1

### Figure S4

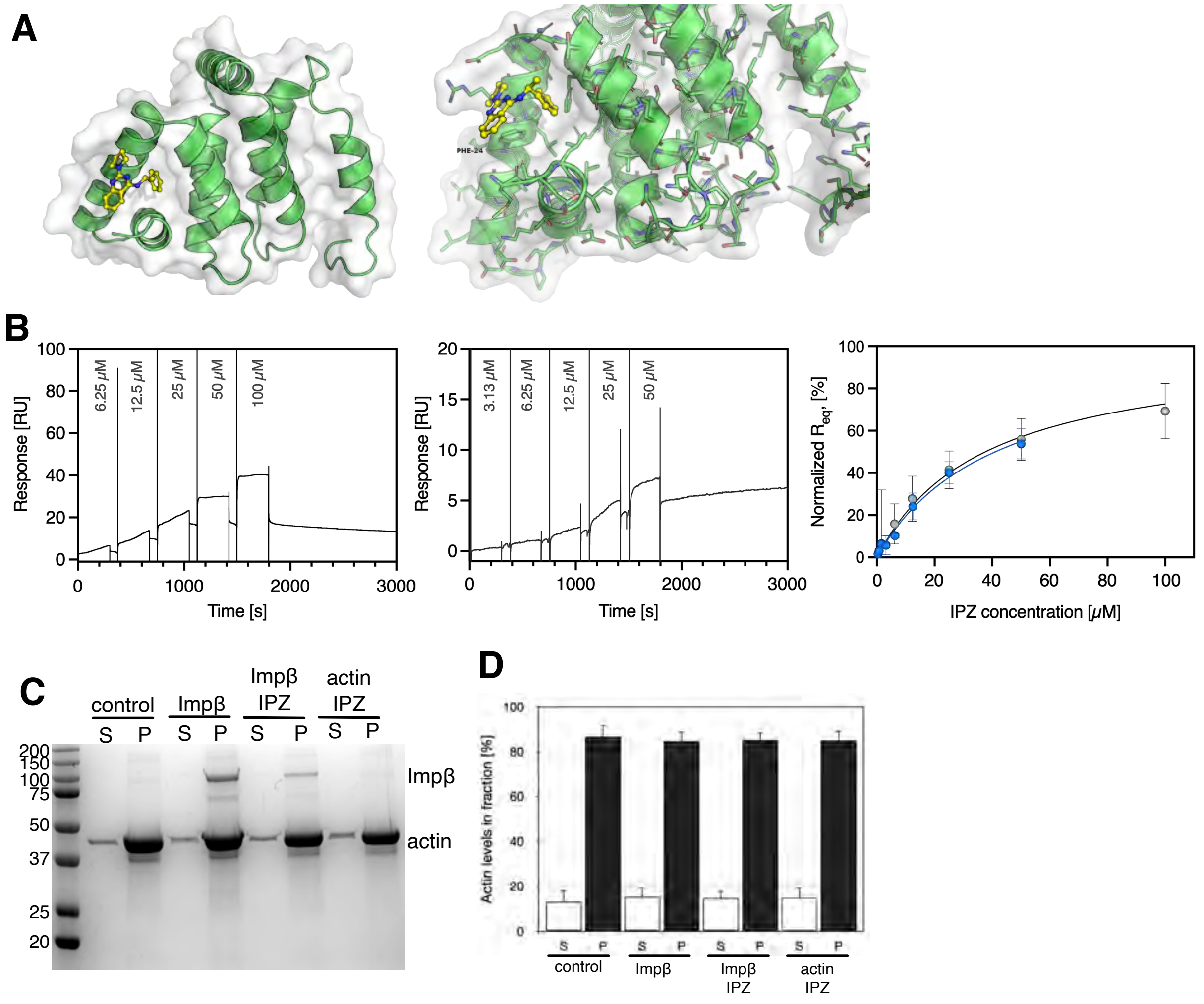
A
PHE-24
B
Response [RU]
6.25 μM
12.5 μM
25 μM
50 μM
100 μM
Time [s]
3.13 μM
Normalized $R_{eq}$, [%]
IPZ concentration [μM]
C
control
Impβ
Impβ IPZ
actin IPZ
S
P
Impβ
actin
D
Actin levels in fraction [%]

### Figure S5

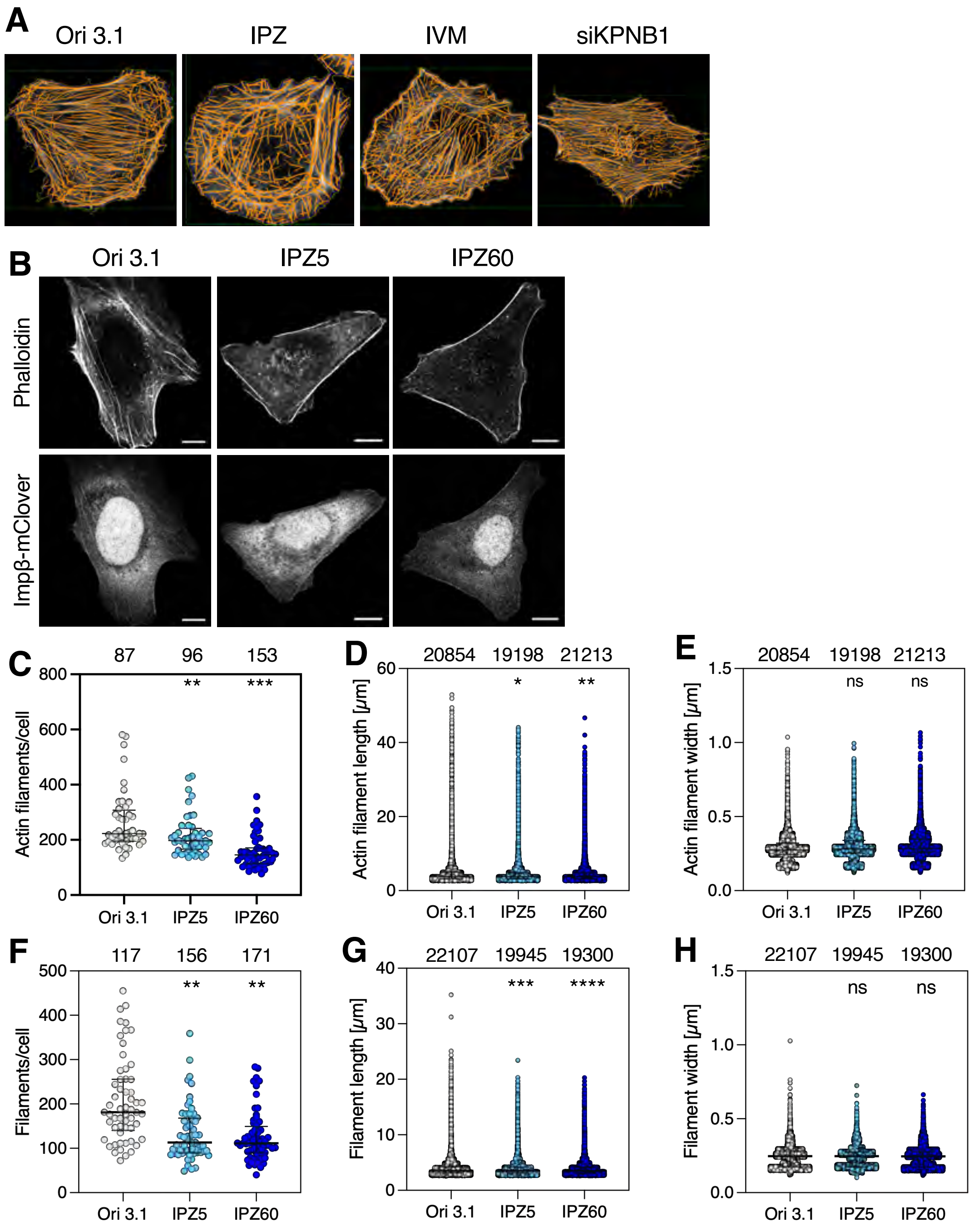
A
Ori 3.1
IPZ
IVM
siKPNB1
B
Ori 3.1
IPZ5
IPZ60
Phalloidin
Impβ-mClover
C
87
96
153
**
***
Actin filaments/cell
800
600
400
200
0
Ori 3.1
IPZ5
IPZ60
D
20854
19198
21213
*
**
Actin filament length [μm]
60
40
20
0
Ori 3.1
IPZ5
IPZ60
E
20854
19198
21213
ns
ns
Actin filament width [μm]
1.5
1.0
0.5
0.0
Ori 3.1
IPZ5
IPZ60
F
117
156
171
**
**
Filaments/cell
500
400
300
200
100
0
Ori 3.1
IPZ5
IPZ60
G
22107
19945
19300
***
****
Filament length [μm]
40
30
20
10
0
Ori 3.1
IPZ5
IPZ60
H
22107
19945
19300
ns
ns
Filament width [μm]
1.5
1.0
0.5
0.0
Ori 3.1
IPZ5
IPZ60

### Figure S6

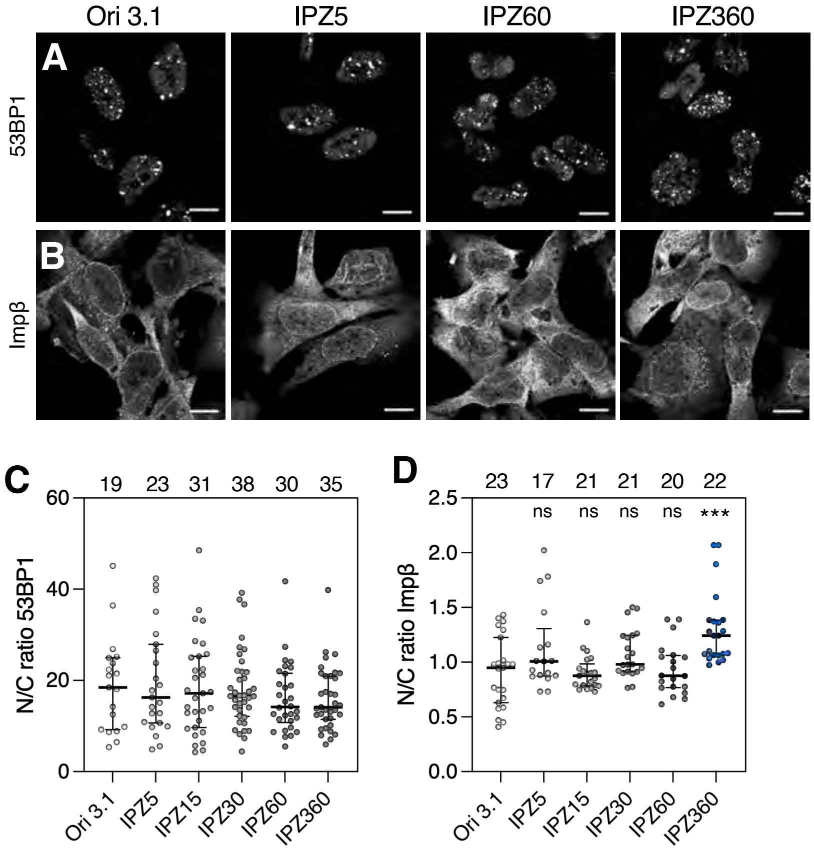
Ori 3.1
IPZ5
IPZ60
IPZ360
A
53BP1
B
Impβ
C
19 23 31 38 30 35
N/C ratio 53BP1
60
40
20
0
Ori 3.1
IPZ5
IPZ15
IPZ30
IPZ60
IPZ360
D
23 17 21 21 20 22
ns ns ns ns ***
N/C ratio Impβ
2.5
2.0
1.5
1.0
0.5
0.0
Ori 3.1
IPZ5
IPZ15
IPZ30
IPZ60
IPZ360

### Figure S7

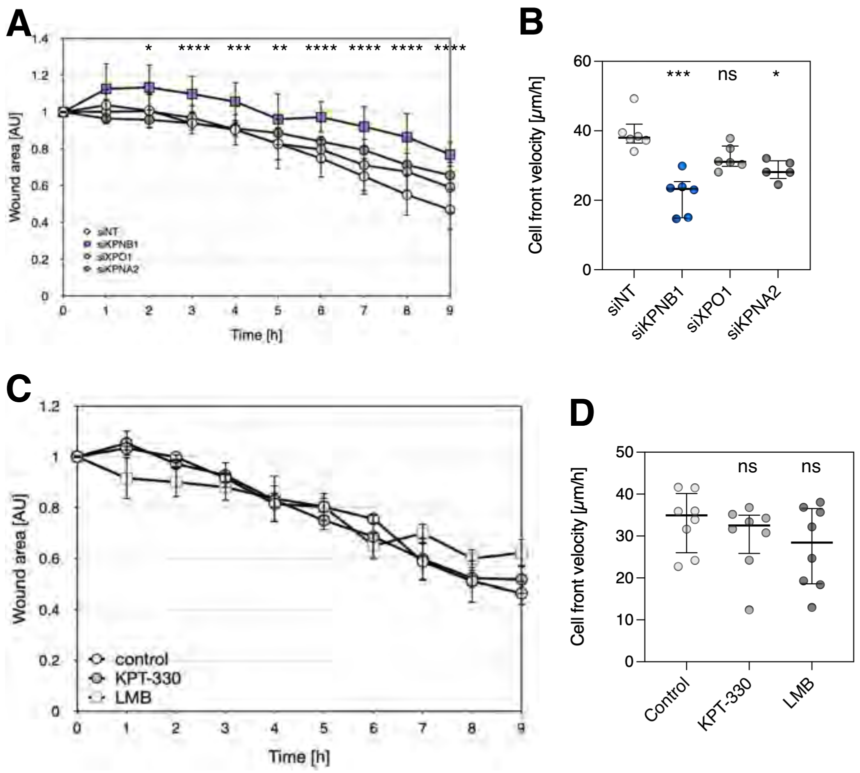
A
Wound area [AU]
Time [h]
siNT
siKPNB1
siXPO1
siKPNA2
* **** *** ** **** **** **** ****
B
Cell front velocity [µm/h]
*** ns *
siNT
siKPNB1
siXPO1
siKPNA2
C
Wound area [AU]
Time [h]
control
KPT-330
LMB
D
Cell front velocity [µm/h]
ns ns
Control
KPT-330
LMB

Supplemental Table S1: Plasmids used in this study

| **Plasmid** | **Construct** | **Source** |
|---|---|---|
| PBF094 | pGEX-6P | Clontech |
| PBF120 | pMK289-g1-KPNB1 | This study |
| PBF121 | pMK392-g1-KPNB1 | This study |
| PBF992 | pDEST-ORF-V1 | Addgene #73637 |
| PBF993 | pDEST-ORF-V2 | Addgene #73638 |
| PRL339 | pX330-U6-Chimeric_NN-CBh-hSpCas9 | Addgene #42230 |
| PRL345 | pMK289 (mAID-mClover-NeoR) | Addgene#72827 |
| PRL405 | mOrange-vinculin-23 | Addgene #57978 |
| PRL472 | Dendra2-actin C-18 | Addgene #57701 |
| PRL502 | mCherry-Cdc42_C-10 | Addgene #55014 |
| PRL584 | mCherry-Arp2-N-14 | Addgene #54980 |
| PRL491 | pDEST-vinculin-V2 | Jühlen and Fahrenkrog, 2023 |
| PRL492 | pDEST-actin-V2 | Jühlen and Fahrenkrog, 2023 |
| PRL661 | pDEST-actin E167A-V2 | Jühlen and Fahrenkrog, 2023 |
| PRL662 | pDEST-actin S350A-V2 | Jühlen and Fahrenkrog, 2023 |
| PRL84 | pKPNB1-EGFP | Patrizia Lavia; (JCB, 2012) |
| PRL720 | pMRX-IPU-GFP-MYH9 | Addgene #168273 |
| PRL489 | pDEST-KPNB1-V1 | This study |
| PRL534 | pDEST-actin-V1 | This study |
| PRL541 | pDEST-KPNB1-Isoform 2-V1 | This study |
| PRL559 | pDEST-α-tubulin-V2 | This study |
| PRL562 | pDEST-KPNB1(1-145)-V1 | This study |
| PRL575 | pDEST-KPNB1(1-145Δ2-31)-V1 | This study |
| PRL589 | pDEST-ARP2-V2 | This study |
| PRL597 | pDEST-KPNB1(1-31)-V1 | This study |
| PRL655 | pDEST-KPNB1 E8A-V1 | This study |
| PRL656 | pDEST-KPNB1 Q22A-V1 | This study |
| RRL657 | pDEST-KPNB1 E8AQ22A-V1 | This study |
| PRL663 | pDEST-actin E167AS350A-V2 | This study |
| PRL710 | pDEST-XPO1-V1 | This study |
| PRL512 | pGEX- KPNB1(1-145) | This study |
| PRL602 | pGEX- KPNB1(1-31) | This study |
| PRL603 | pGEX- KPNB1(1-145Δ2-31) | This study |
| PRL605 | pKPNB1(1-145)-EGFP | This study |

| | | |
|---|---|---|
| PRL606 | pKPNB1(1-145Δ2-31)-EGFP | This study |
| PRL608 | pKPNB1(1-31)-EGFP | This study |
| PRL724 | pKPNB1(Δ2-31)-EGFP | This study |

**Supplemental Table S2**: Primers used in this study

| Primer name | Sequence | Target plasmid |
|---|---|---|
| Vector V1 For | tttctcgttcagctttcttgtacaaagtg | pDEST-KPNB1-V1 |
| Vector V1 Rev | gcttttttgtacaaacttgtctcgagtacc | |
| V1-ImpB For | tttgtacaaaaaagcatggagctgatcaccattctcg | |
| V1-ImpB Rev | aagctgaacgagaaaTTAcgcttggttcttcagtttcctca | |
| ImpB.For | gaaactgaagaaccaagcgtcgAACCCAGCTTTcttgtac | |
| ImpB.Rev | gtacaagAAAGCTGGGTTcgacgcttggttcttcagtttc | |
| V1/V2 For | AACCCAGCTTTcttgtacaaagtgg | pDEST-ORF-V1, pDEST-ORF-V2 |
| V1/V2 Rev | catTAAGCCTGCTTTTTTGTACaaacttgtct | |
| V1 Actin FOR | AAAGCAGGCTTAatggatgatgatatcg | pDEST-actin-V1 |
| V1 Actin REV | caagAAAGCTGGGTTgaagcatttgcggt | pDEST-actin-V1 |
| ImpB-145 For | ATGAAGGAGTCGACATTGGAAG | pDEST-KPNB1-Isoform 2-V1 |
| V1-ImpB -145 Rev | TAAGCCTGCTTTTTTGTAC | |
| Tubulin FOR | AAAGCAGGCTTAATGCGTGAGTGCATCT | pDEST-α-tubulin-V2 |
| Tubulin REV | CAAGAAAGCTGGGTTGTATTCCTCTCCTTCTTC CTCACCC | pDEST-α-tubulin-V2 |
| V1-N Term ImpB For | caacagcacagagcacAACCCAGCTTTctt | pDEST-KPNB1(1-145)-V1 |
| V1-N Term ImpB Rev | gtgctctgtgctgttggggtttg | |
| Ran Insert.For | AAAAAAGCAGGCTTAATGCCTGCGCAGGG | pDEST-RAN-V1 |
| RanInsert.Rev | caagAAAGCTGGGTTCAGGTCATCATCCTCATCCGGG | |
| Imp 2-31 For | AAAGCAGGCTTAatgaacctgcccactttccttg | pDEST-KPNB1(1-145Δ2-31)-V1 |
| Imp 2-31 Rev | catTAAGCCTGCTTTTTTGTAC | |
| Fragm ARP For | cAAAAAAGCAGGCTTAatggacagccag | pDEST-ARP2-V2 |
| Fragm ARP Rev | CAAGAAAGCTGGGTTtcgaacagtcacaccaagtttctct | |
| V1-ImpB 1-31 For | cgtgcggccgtggagAACCCAGCTTTcttgtacaaag | pDEST-KPNB1(1-31)-V1 |
| V1-ImpB 1-31 Rev | ctccacggccgcacgctccag | |
| Fragm RhoA For | AAAGCAGGCTTAatggctgccatccggaagaaactg | pDEST-RhoA-V2 |
| Fragm RhoA Rev | caagAAAGCTGGGTTCAAGACAAGgcaaccagattt | |
| Fragm Rac1 For | AAAGCAGGCTTAatgcaggccatcaagtgtgtggtg | pDEST-Rac1-V2 |
| Fragm Rac1 Rev | caagAAAGCTGGGTTcaacagcaggcattttctcttcct | |
| V1-XPO1 Insert FOR | AAAGCAGGCTTAATGCCAGCAATTATGACAATGTTAGC | pDEST-XPO1-V1 |
| V1-XPO1 Insert Rev | CAAGAAAGCTGGGTTATCACACATTTCTTCTGGAATCTCATGT | |
| pGex-ImpB del 2-31 For | CCAGCGGCCGCCATGAACCTGCCCACTTTCC | pGEX- KPNB1(1-145Δ2-31) |
| pGex-ImpB del 2-31 Rev | CATGGCGGCCGCTGGAACAG | |
| pEGFP-Impb1-145 For | caacagcacagagcacgatccaccggtcgccaccatg | pKPNB1(1-145)-EGFP |
| pEGFP-Impb1-145 Rev | gtgctctgtgctgttgggg | |
| pEGFP-ImpB del 2-31 For | gatctcgagctcatgaacctgcccactttccttgtg | pKPNB1(1-145Δ2-31)-EGFP |
| pEGFP-ImpB del 2-31 Rev | atgagctcgagatctgagtc | |

| | | |
|---|---|---|
| pEGFP-Impb1-31 For | cgtgcggccgtggaggatccaccggtcgccaccatg | pKPNB1(1-31)-EGFP |
| pEGFP-Impb1-31 Rev | ctccacggccgcacgctccag | |
| pEGFP-ImpB del 2-31 For | gatctcgagctcatgaacctgcccactttccttgtg | pKPNB1(Δ2-31)-EGFP |
| pEGFP-ImpB del 2-31 Rev | atgagctcgagatctgagtc | |
| ImpB A8 For | atggagctgatcaccattctcgcgaagaccgtgtctcccgatcg | pDEST-KPNB1 E8A-V1 |
| ImpB A8 Rev | cgatcgggagacacggtcttcgcgagaatggtgatcagctccat | |
| ImpB A22 For | ctggagctggaagcggcggcgaagttcctggagcgtgcg | pDEST-KPNB1 Q22A-V1 |
| ImpB A22 Rev | cgcacgctccaggaacttcgccgccgcttccagctccag | |
| Actin A167 For | cactgtgcccatctacgcggggtatgccctcccccatg | pDEST-actin E167AS350A-V2 |
| Actin A167 For | catgggggagggcataccccgcgtagatgggcacagtg | |
| Actin A350 For | gctccatcctggcctcgctggccaccttccagcagatgtg | |
| Actin A350 Rev | cacatctgctggaaggtggccagcgaggccaggatggagc | |
| ImpB 145 Stop For | CCAACAGCACAGAGCACTGAAGGAGTCGACATTG | pGEX- KPNB1(1-145) |
| ImpB 145 Stop Rev | CAATGTCGACTCCTTCAGTGCTCTGTGCTGTTGG | |
| ImpB 31 Stop For | GAGCGTGCGGCCGTGGAGtaaCTGCCCACTTTCCTTGTGG | pGEX- KPNB1(1-31) |
| ImpB 31 Stop Rev | CCACAAGGAAAGTGGGCAGttaCTCCACGGCCGCACGCTC | |
